# Genetic Drift and Selection in Many-Allele Range Expansions

**DOI:** 10.1101/145631

**Authors:** Bryan T. Weinstein, Maxim O. Lavrentovich, Wolfram Möbius, Andrew W. Murray, David R. Nelson

## Abstract

We experimentally and numerically investigate the evolutionary dynamics of four competing strains of *E. coli* with differing expansion velocities in radially expanding colonies. We compare experimental measurements of the average fraction, correlation functions between strains, and the relative rates of genetic domain wall annihilations and coalescences to simulations modeling the population as a one-dimensional ring of annihilating and coalescing random walkers with deterministic biases due to selection. The simulations reveal that the evolutionary dynamics can be collapsed onto master curves governed by three essential parameters: (1) an expansion length beyond which selection dominates over genetic drift; (2) a characteristic angular correlation describing the size of genetic domains; and (3) a dimensionless constant quantifying the interplay between a colony’s curvature at the frontier and its selection length scale. We measure these parameters with a new technique that precisely measures small selective differences between spatially competing strains and show that our simulations accurately predict the dynamics with no additional fitting. Our results suggest that that the random walk model can act as a useful predictive tool when describing the evolutionary dynamics of range expansions composed of an arbitrary number of competing alleles with different fitnesses.

**Author summary:** Population expansions occur naturally during the spread of invasive species and have played a role in our evolutionary history when many of our ancestors expanded out of Africa. We use a colony of bacteria expanding into unoccupied, nutrient-rich territory on an agar plate as a model system to explore how an expanding population’s spatial structure impacts its evolutionary dynamics. Spatial structure is present in expanding microbial colonies because daughter cells migrate only a small distance away from their mothers each generation. Generally, the constituents of expansions occurring in nature and in the lab have different genetic compositions (genotypes, or alleles if a single gene differs), each instilling different fitnesses, which compete to proliferate at the frontier. Here, we show that a random-walk model can accurately predict the dynamics of four expanding strains of *E. coli* with different fitnesses; each strain represents a competing allele. Our results can be extended to describe any number of competing genotypes with different fitnesses in a naturally occurring expansions. Our model can also be used to precisely measure small selective differences between spatially competing genotypes in controlled laboratory settings.

## Introduction

A competition between stochastic and deterministic effects underlies evolution. In a well-mixed system such as a shaken culture of the yeast microorganism *Saccharomyces cerevisiae*, stochastic competition between individuals, mutations, and selection dictate the dynamics of the population [1]. In structured (i.e. spatial) environments, active or passive dispersal of individuals also plays an important role. The local “well-mixed” dynamics must be coupled to the motion of individuals, leading to strikingly different global population dynamics, even in the absence of selection [2–7].

A model laboratory system that can be used to explore the coupling between local “well-mixed” effects and spatial deterministic and stochastic dynamics is a microbial range expansion [8], in which a population expands into an unoccupied region of a hard agar Petri dish. Non-motile microbes expand outwards from their initial position due to a combination of growth coupled with random pushing by neighboring cells and leave behind a record of their genetic competition as they cannot move and cease reproducing once the population becomes too dense [8]. A frozen genetic pattern of four competing strains of *E. coli* marked by different fluorescent colors can be seen in Figure 1. Spatial structure is present in the frozen genetic patterns because the microbes at the expanding frontier produce daughter cells of the same color that migrate only a small fraction of the front circumference within a generation. Hallatschek et al. [8] identified the key role of genetic drift in producing these sectored patterns; the small population size at the front of an expanding population [9, 10] enhances number fluctuations (i.e. genetic drift), eventually leading to the local fixation of one strain past a critical expansion radius *R*_0_. The decrease in genetic diversity as the small number of individuals at the frontier expands is referred to as the “Founder effect” [11].

**Fig 1.**
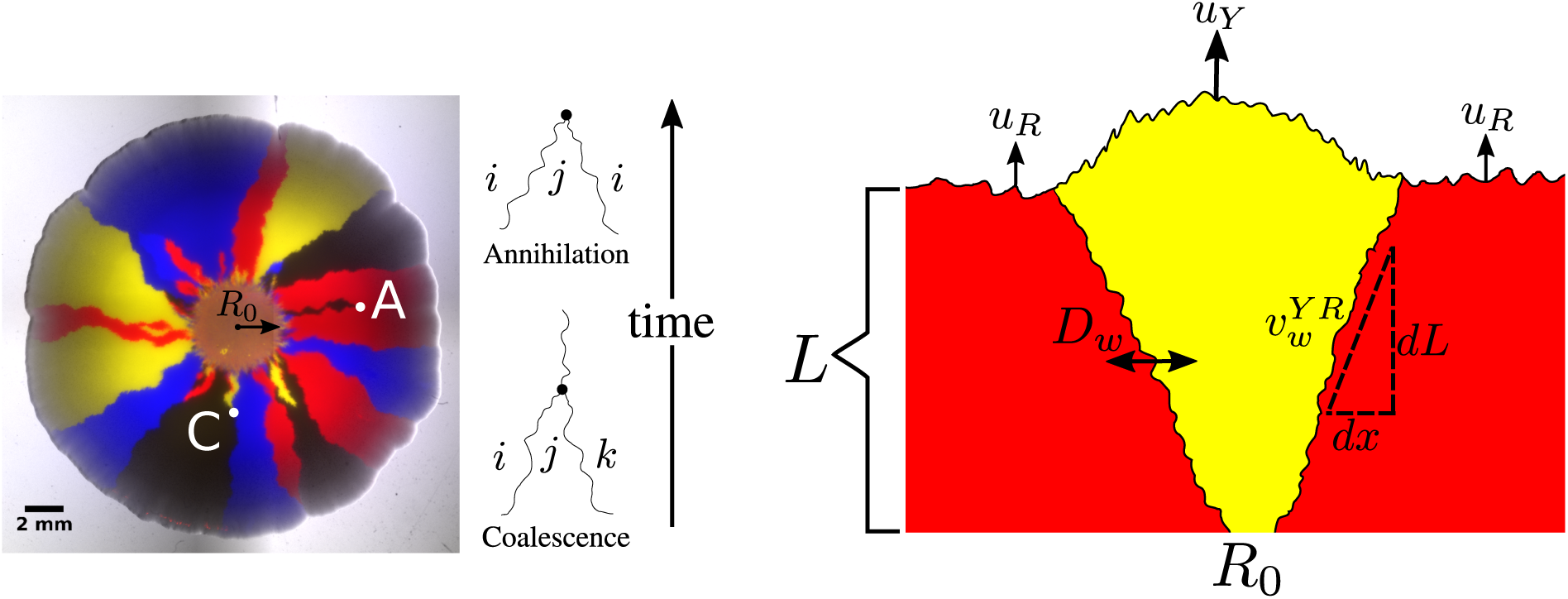
*Left*: A four-color *E. coli* range expansion. Four strains of *E. coli* differing only by a heritable fluorescent marker were inoculated on an agar plate in a well-mixed droplet and expanded outwards, leaving behind a “frozen record” of their expansion. Virtually all growth occurred at the edge of the colony. The markers instilled different expansion velocities: our eCFP (blue) and eYFP (yellow) strains expanded the fastest, followed by our black strain, and finally our mCherry (red) strain. As a result of the differing expansion velocities, the yellow/blue bulges at the frontier are larger than the black bulges which are larger than the red bulges, although the significant stochastic undulations at the front mask their size. The microbes segregate into one color locally at a critical expansion radius *R*_0_ due to extreme genetic drift at the frontier [8]. After segregated domains form, genetic domain walls diffuse and collide with neighboring walls in an “annihilation” or “coalescence” event indicated by an *A* or *C* respectively. *Right:* A faster expanding, more fit yellow strain with expansion velocity *u_Y_* sweeping through a less fit red strain with expansion velocity *u_R_* in a regime where the curvature of the colony can be neglected. Throughout this paper, we quantify the dynamics of the sector as a function of length expanded by the colony, *L* = *R* − *R*_0_. We characterize domain wall motion per differential length expanded (the radius the colony has grown) *dL* and the wall’s differential displacement perpendicular to the expansion direction *dx*. 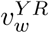 is the yellow-red (*Y R*) domain wall’s average selection-induced deterministic displacement *dx* perpendicular to the expansion direction per length expanded *dL*. *D_w_* is the domain walls’ diffusion coefficient per length expanded; it controls how randomly the domain walls move. We model the dynamics of our four strains as a one-dimensional line of annihilating and coalescing random walkers using the parameters *R*_0_, *D_w_*, and 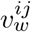, where *ij* represents all possible types of domain walls.

Outside of the laboratory, range expansions occur naturally during the spread of invasive species such as the bank vole in Ireland [12] or the cane toad in Australia [13], and played a role in the evolutionary history of humans when many of our ancestors expanded out of Africa [14]. In these natural expansions, populations may have many competing genotypes, or alleles, each instilling a different fitness. Even if a population is originally clonal, mutations may create new alleles that compete with one another to proliferate, a phenomenon known as clonal interference [15].

An allele’s fitness is often determined by its corresponding expansion velocity. Faster expanding individuals will colonize more territory and will block slower strains from expanding, resulting in the increased abundance of faster alleles at the frontier [13, 16, 17]. If the curvature of a microbial colony can be neglected and its front is sufficiently smooth, it has been shown both theoretically and experimentally that the domain wall of a faster expanding strain will displace a slower expanding strain at a constant rate per length expanded, resulting in a characteristic triangular shape [17] as shown on the right side of Figure 1. If the curvature of the expansion is not negligible, the sector boundaries will trace logarithmic spirals [17].

Even in the most simple scenario when de-novo mutations and mutualistic or antagonistic interactions are ignored, the dynamics of *many* competing alleles with varying fitnesses at the front of a range expansion have not been quantified theoretically nor explored in laboratory experiments. Prior laboratory experiments focused on the dynamics of a single sector of a more fit strain (representing a competing alelle) of yeast or bacteria sweeping through a less fit strain [17] in regimes where stochastic wandering of genetic boundaries was not expected to be important. Recent experimental work studied how fast a *single* more fit strain swept through a less fit strain in a range expansion and compared the dynamics to the same strains in a well mixed test tube [9].

In this paper, we experimentally and numerically investigate the dynamics of four competing strains (alleles) of *E. coli* with varying selective advantages initially distributed randomly at the front of a radial range expansion. The eCFP (blue) and eYFP-labeled (yellow) strains expanded the fastest, followed by the non-fluorescent (black) strain, and finally the mCherry-labeled (red) strain. The differences in expansion speeds are reflected in Figure 1 as follows: the yellow/blue bulges at the front of the expansion are larger than the black bulges which are larger than the red bulges. The significant random undulations at the frontier, however, significantly mask the selection-induced bulges.

As is evident from Figure 1, the size and location of a monoclonal sector can be described by the locations of its boundaries. We therefore describe our expansions as a one-dimensional line of annihilating and coalescing random walkers, a description that has been used extensively in previous work (see Ref. [2] for a review). To account for the radial geometry of our colonies, we allow the frontier to inflate, corresponding to the increasing perimeter of the colony as its radius increases. Past the radius *R*_0_ where genetic domains originally form, we describe the random motion of genetic domains by a diffusion constant per length expanded *D_w_* (see Figure 1) [18]. If *dx* characterizes the displacement of a domain wall perpendicular to the expansion direction and *dL* is the distance the colony has expanded (the radius that the colony has grown) as illustrated on the right side of Figure 1, where we neglect the circumferential curvature in this small region, we define the diffusion constant per length expanded as 2*D_w_* = *d*Var(*x*) / *dL* where Var(*x*) ≡ 〈*x*^2^〉 − 〈*x*〉^2^ is the variance and the brackets indicate an average over many domain walls. Note that *D_w_* has dimensions of length. Similarly, differences in expansion velocities between neighboring strains will lead to the deterministic displacement of domain walls per length expanded as the faster expanding strain will reach the contested point on the front before a slower growing strain as mentioned above [17]; we characterize this deterministic motion by a dimensionless “wall velocity,” [18] 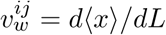, where *i* is the strain to the left of the domain wall and *j* is the strain to the right. Note that 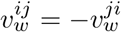.

The dynamics of an arbitrary number of *neutral* competing strains in an expansion (i.e. 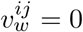 for all domain walls) is well understood as the dynamics can be described as a one-dimensional system of annihilating and coalescing random walkers [19–21] which is equivalent to a one-dimensional *q*-state Potts model [22, 23] governed by zero-temperature Glauber dynamics [24] or a *q*-opinion Voter model [25, 26]. Many theoretical predictions and analyses of this system exist; of particular relevance to this paper are the relative annihilation and coalescence rates per collision as *q* is varied [27–29] and the calculation of spatial correlation functions [28]. To map standard linear results onto an inflating ring (i.e. including *R*_0_ in the models), one can use a conformal time transformation [30–32]. Fewer results are available in the presence of selection, i.e. when domain walls have nonzero 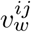 [33]. Analytical results are rare because the moment hierarchy of this model does not close [2] as discussed in Appendix S1.

In this paper, we measure and predict three quantities: the average fraction of each of our four strains, the two-point correlation functions between our strains, and the relative annihilation and coalescence probabilities per domain wall collision (see Figure 1), a quantity that has received theoretical attention [27–29] but has not been explored experimentally nor investigated in the presence of selection. We measure these three quantities using an image analysis toolkit (available on Github [34]) that extends experimental techniques for two-color (two-allele) range expansions [8, 9, 17, 18, 35, 36] to an arbitrary number of competing strains. We next use an efficient radial simulation (also on Github [34]) of annihilating and coalescing random walkers with deterministic wall velocities to determine what sets the scale of the dynamics and to synthesize our experimental and theoretical results. We show that key combinations of *R*_0_, *D_w_* and 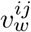 control the dynamics of our four strains. We conclude with suggestions for future studies.

## Materials and methods

### Experimental setup

We used four DH5*α* strains of *E. coli* (labeled BW001, BW002, BW003, and BW012), each expressing spectrally distinguishable fluorescent colors. The unique colors were obtained by using the plasmid vector pTrc99a [37] with an open reading frame for the respective fluorescent proteins. Strains BW001, BW002, and BW003 expressed eCFP (cyan/blue), eYFP (yellow), and mCherry (red) respectively, and were identical to the *E. coli* strains eWM282, eWM284, and eWM40 used in ref. [38]. Strain BW012 was a mutated descendant of strain BW002 (yellow) that fluoresced at a decreased intensity, appearing black, while retaining its ampicillin resistance from the pTrc99a vector. Throughout this work, no additional mutations were introduced or observed. We therefore consider that these four strains correspond to four different alleles. Throughout the rest of the paper, we refer to the strains as eCFP, eYFP, mCherry, and black.

To prepare saturated cultures, strains were inoculated in 10mL of 2xYT media and were shaken for approximately 16 hours at 37°C. After vortexing each saturated culture and obtaining their concentration via optical density (OD-600) measurements, appropriate volumes (e.g., 1:1:1 mixtures of three strains) were added to an Eppendorf tube with a final volume of 1mL. The Eppendorf tube was then vortexed to uniformly mix the strains. A volume of 2 *μ*L was taken from the vortexed tube and placed on center of a 100 mm diameter Petri dish containing 35 mL of a lysogeny broth (LB), ampicillin, and 1.25% w/v bacto-agar. The carrier fluid in the resulting circular drop evaporated within 2-3 minutes, depositing a circular “homeland” of well-mixed bacteria onto the plate.

After inoculation, plates were stored for 8 days upside down (to avoid condensation) in a Rubbermaid 7J77 box at 37°C with a beaker filled with water; the water acted as a humidifier and prevented the plates from drying. The plates were occasionally removed from the box and imaged (at roughly 24 hour intervals) using the brightfield channel to determine the radius of the colony as a function of time. On the eighth day, the plates were imaged in both fluorescent and brightfield channels. The number of replicate plates used are stated below next to the respective experimental results.

### Image acquisition and analysis

We imaged our range expansions with a Zeiss SteREO Lumar.V12 stereoscope in four channels: eCFP, eYFP, mCherry, and brightfield. In order to analyze a colony with a maximum radius of approximately 10 mm using a single image, we stitched four images together with an overlap of 20% using AxioVision 4.8.2, the software accompanying the microscope. We blended the overlapping areas of the images to lessen the impact of background inhomogeneities. An example of a stitched image can be seen on the left side of Figure 2. Stitching introduced small artifacts such as vertical lines near the center of our expansions; we verified that these did not affect our results.

**Fig 2.**
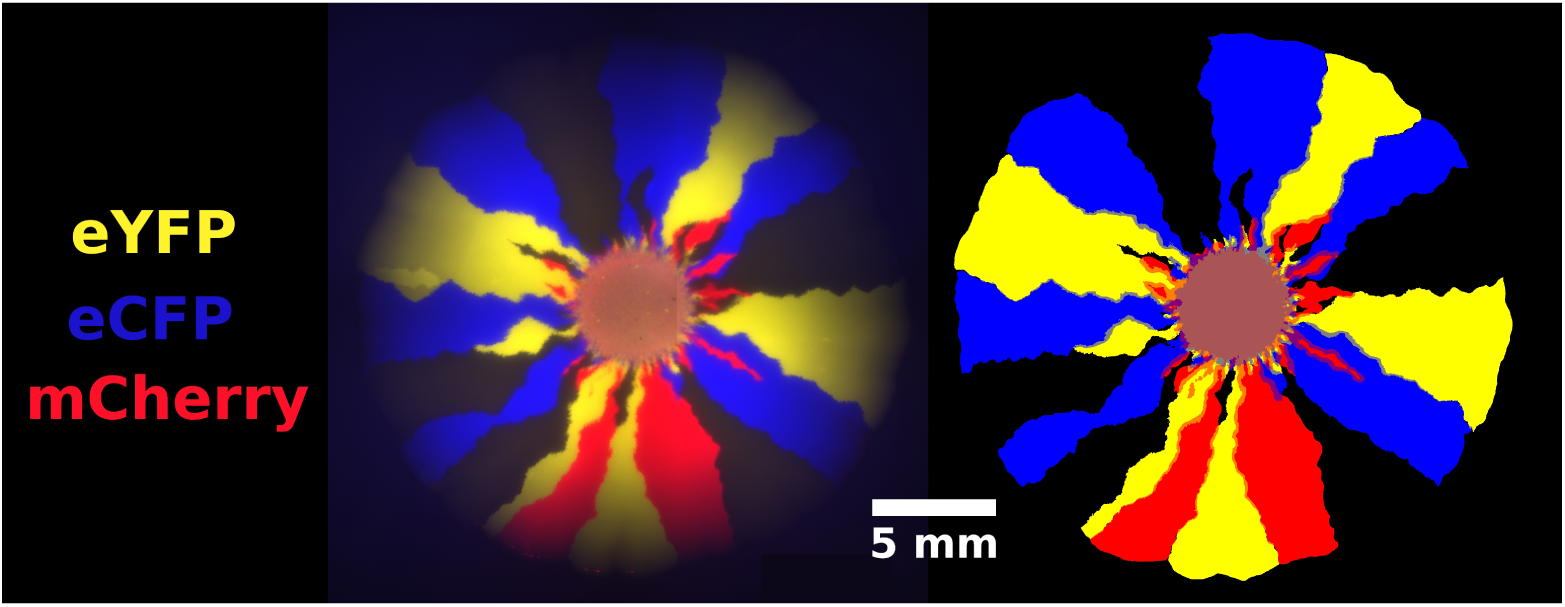
A four-color *E. coli* range expansion (left) and the superimposed binary masks of each channel (right) [34]. Images were acquired for four overlapping quadrants and stitched together to obtain a single image with a large field of view. Overlapping regions were blended to minimize inhomogeneities. To obtain the binary masks, pixels with fluorescence above background noise were marked as “on.” A visual comparison of the raw data and the masks confirm that our binary masks accurately reflect the location and shape of individual sectors.

To extract the local fraction of each strain per pixel, we first created binary masks for each fluorescence channel indicating if the corresponding *E. coli* strain was present. We utilized the “Enhance Local Contrast” (CLAHE) algorithm [39] in Fiji [40], an open-source image analysis platform, to help correct for inhomogeneities in background illumination. After applying the CLAHE algorithm, a combination of automatic thresholding and manual tracing yielded a binary mask of each channel, an example of which is shown in Figure 2; the image on the left is an overlay of an experimental range expansion’s fluorescent channels and the image on the right is the overlay of the corresponding binary masks.

We mapped the binary images to the local fraction of each *E. coli* strain in the following way : if *N* binary masks (corresponding to *N* colors) were “on” at a pixel, the local fraction of their corresponding channels was assigned to be 1/*N*. Although this assignment produces inaccuracies (i.e., if one strain occupied 90% of a pixel and the other occupied 10%, our algorithm would register both as 50%), domain boundaries were the only areas besides the homeland and the early stages of the range expansions where multiple strains were colocalized. The black strain (with an inactivated fluorescent protein) was defined to be present at pixels reached by the range expansion in which no other strains were present. Although this definition introduced errors at radii close to the homeland with significant color overlap, the error became negligible at large radii as quantified in Supplementary Figure S1. Once we determined the fraction of each strain at each pixel, we were able to extract quantities such as the total fraction of each strain in the colony and spatial correlations between strains at a given expansion radius.

The mask in Figure 2 highlights that sector boundaries can be used to determine local strain abundance. Although it is possible to extract the position of every domain wall from each strains’ local fraction, it is challenging to actually *track* a single wall due to collisions between walls. To address this problem, we created a binary mask of the edges in our images and labelled the edges of each domain. Annihilations and coalescences were counted manually within Fiji [40]; automated measures were not accurate enough.

### Characterizing radial expansion velocities

We used the average expansion velocity of each strain for radii *R* > *R*_0_ as a proxy for selective advantage, similar to previous work [17, 35]. We measured the diameter of 12 expansions approximately every 24 hours over the course of 7 days. To account for biological variance, every four of the 12 colonies were created from independent single colonies; no statistical difference was seen between biological replicates. The diameters were determined by manually fitting a circle to a brightfield image of the expansion three times and averaging the measured diameters. Figure 3 shows the average radius increasing with time for each strain. The eCFP and eYFP strains had the fastest expansion velocities (the respective datapoints overlap in Fig. 3), followed by the black strain, and then finally the mCherry strain. The expansion velocity slowly decreased as a function of time, which we attribute to nutrient depletion in the plate.

**Fig 3.**
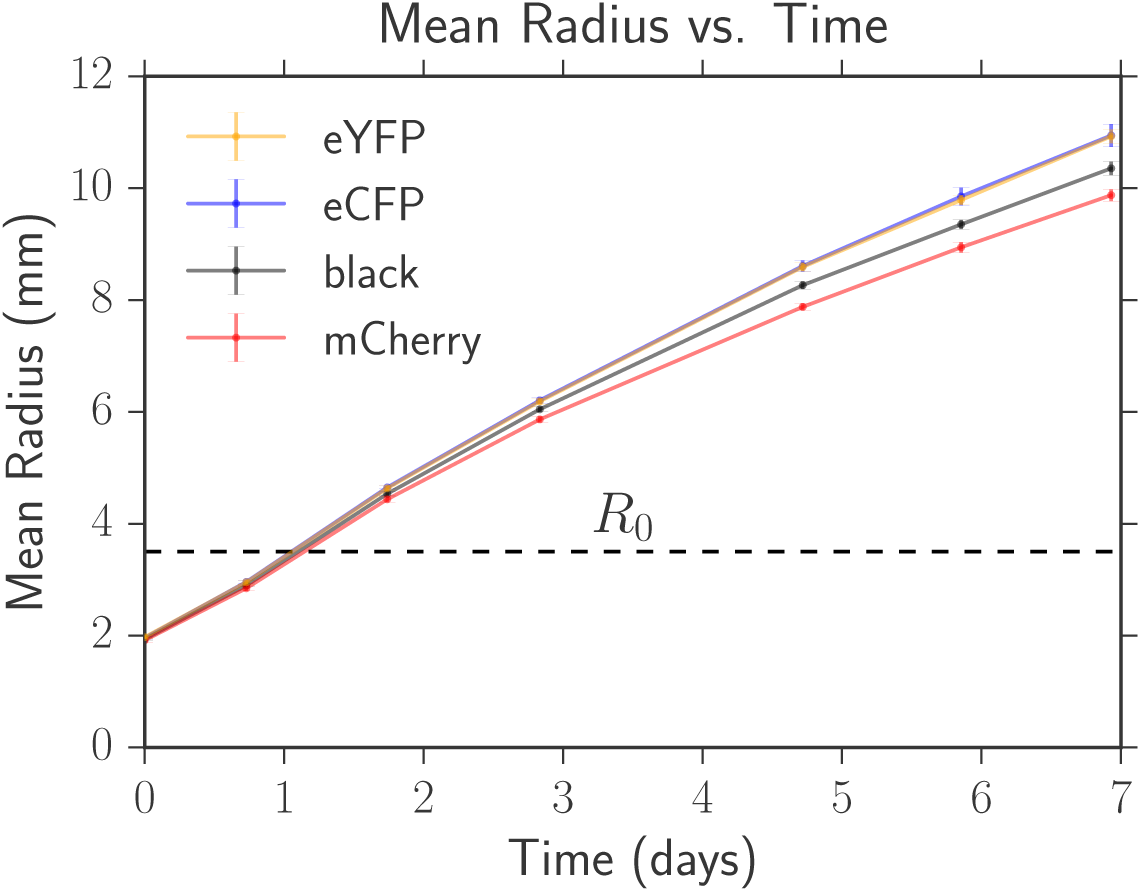
The average radius versus time for each strain. The error bars (comparable to symbol size at early times) are the standard errors of the mean calculated from 12 replicate expansions for each strain. The eYFP and eCFP strains had the fastest expansion velocities (data points overlap in the plot) followed by black and then mCherry. *R*_0_ is the radius at which expansions with competing strains typically demix into one color locally; *R*_0_ is approximately 1.75 times the initial inoculant radius of 2 mm (see Fig. 1).

The average radial expansion velocity of each strain was obtained by using linear regression to fit the radius versus time for radii greater than *R*_0_. To estimate the error in the expansion velocities we followed a bootstrapping approach; we sampled with replacement from each of the 12 radii trajectories, and fit a slope to the average radius versus time of the sampled data. The mean and error were the mean and standard deviation of the sampled *u_i_* distributions for each strain *i*. Our calculated values of *u_i_* appear in Table 1.

We then quantified the dimensionless selective advantage of each strain relative to the slowest growing mCherry strain following [17] via *s_iR_* = *u_i_*/*u_R_* − 1 where the *R* indicates the mCherry strain (red); see Table 1. The selective advantages were consistent, within error, when we calculated the velocities *u_i_* and *u_R_* over different time intervals. The eCFP and eYFP strains had a selective advantage of 16%, similar to the experiments of Weber et al. [35] which found, despite the fact that they used different *E. coli* strains and plasmids, that the expression of mCherry decreased the expansion velocity of their strains by approximately 15% in certain their “fast growth” environmental conditions. Our black strain had an approximately 8% enhancement over the mCherry strain. When we took the plasmids from our strains and inserted them into clonal dH5*α* strains, we saw the same fitness defects as judged by a decrease in average fraction in competition experiments, indicating that the fitness defect was indeed due to the presence of the plasmids and expression of the corresponding fluorescent proteins. Selective advantages of this magnitude were observed when studying yeast *S. cerevisiae* range expansions [17].

**Table 1.** The average expansion velocity *u_i_* and each strain’s selective advantage relative to the mCherry strain *s_iR_* = *u_i_*/*u_R_* − 1 measured over the course of seven days for radii greater than *R*_0_ (the radius where distinguishable domain walls formed).

| Strain | Average Velocity $u_i$ (mm/day) | $s_{iR} = u_i/u_R - 1$ |
| --- | --- | --- |
| eYFP | $1.21 \pm 0.02$ | $0.16 \pm 0.03$ |
| eCFP | $1.22 \pm 0.04$ | $0.16 \pm 0.04$ |
| Black | $1.12 \pm 0.02$ | $0.08 \pm 0.02$ |
| mCherry | $1.04 \pm 0.02$ | 0 |

Between sets of plates made on different days, the average radial expansion velocity of each strain changed slightly (at most a deviation of 10%), but the order of expansion velocities between the strains was consistent; the eCFP and eYFP strains were neutral followed by the black strain and then the red strain. Importantly, although the radial expansion velocities varied between sets of plates, the demixing radius *R*_0_, wall velocities 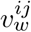, and diffusion coefficient *D_w_* did not within error as discussed in the Quantifying Parameters section, so the evolutionary dynamics of our strains were consistent between sets of plates.

### Simulations

Lattice simulations of range expansions, especially radial ones, can suffer from artifacts arising from the preferred directions of the lattice. It is possible to use an amorphous Bennett model lattice [41] to mitigate some of these effects [32]. Instead, we develop a simple off-lattice method that treats the domain walls as annihilating and coalescing random walkers moving along a continuous line. We incorporate both the random, diffusive motion of the domain walls as well as deterministic movement due to selection, allowing us to simulate the growth of linear (appropriate when the curvature of the colony can be ignored) and radial range expansions. We discuss the linear case first and then extend to the radial case; the linear simulation steps are as follows:

1. Create a line of *N*_0_ microbes of width *a* at the linear frontier. Assign each microbe one of the *q* potential alleles.
2. Identify genetic domain walls by locating neighbors with different alleles; assign type *ij* to each wall where *i* and *j* are the strains to the left and right respectively. Assign a relative “growth rate” *r_ij_* to each wall characterizing the probability that strain *i* divides before strain *j*.
3. Choose a wall at random and move it a distance *a* (the width of the cells) to the left or right; this represents the competition to reproduce and occupy new space at the population frontier (genetic drift). We use periodic boundary conditions for the domain wall positions along the line.

(a) Jump to the right with a probability of 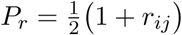 or to the left with probability 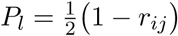. Note that domain walls separating neutral strains (*r_ij_* = 0) will jump to the left or right with equal probability and that *r_ij_* ≤ 1.
(b) If the hopping domain wall collides with another wall, react the walls instantaneously with an appropriate annihilation or coalescence depending on whether the leftmost and rightmost strains are the same or different respectively.
4. Increment the elapsed time by Δ*t* = 1/*N* generations, where *N* is the number of domain walls at the beginning of the jump, and increment the length expanded by the colony by Δ*L* = *d*Δ*t* = *d*/*N*, where *d* is the distance that the colony expands per generation.
5. Repeat steps 3 and 4 until no domain walls remain or until the simulation has run for the desired number of generations.

Our algorithm, thus far simulating only linear expansions, can easily be extended to simulate radial geometries. To incorporate the radially inflating perimeter, we note that a domain wall at a radius *R* will jump an angular distance of *δϕ* = *a*/*R*. As the radius of our experimental expansions increases approximately linearly with generation time, we describe its radius as *R* = *R*_0_ + *d* · *t*, where *t* is measured in generation times. We thus account for inflation by using a time-varying angular jump length of

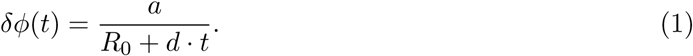

If there are *N*_0_ individuals at the frontier, *R*_0_ is given by *R*_0_ = *N*_0_*a*/(2*π*).

Our simulation’s diffusion coefficient per length expanded, characterizing the random motion of the domain walls, can be shown to be *D_w_* = *a*^2^/(2*d*) while its wall velocity per length expanded, characterizing the deterministic displacement of domain walls due to selection, can be shown to be 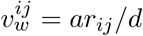. As *r_ij_* ≤ 1, the maximum wall velocity attainable in this simulation is *a*/*d*. In our simulations, it is convenient to set *d* = *a* without loss of generality so that the front of the expansion advances by a single cell width per generation, although *d* could be chosen arbitrarily in the simulation framework. The length expanded is then given by *L* = *a* · *t*, the diffusion coefficient per length expanded by *D_w_* = *a*^2^/(2*d*) = *a*/2, and the wall velocity per length by 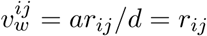.

In contrast to algorithms that follow the position and state of every organism at the front of a colony, our algorithm only tracks the positions of domain walls and is consequently much faster per generation as the sectors coarsen, allowing for simulations of larger colonies. Figure 4 displays a radial and linear simulation with three neutral colors and a fourth red color with a selective disadvantage comparable to our experiments. We check that our simulation correctly reproduces the behavior of a single more fit domain wall sweeping through a less fit strain in Supplementary Figure S5. Our implementation of this algorithm is available on Github [34].

**Fig 4.**
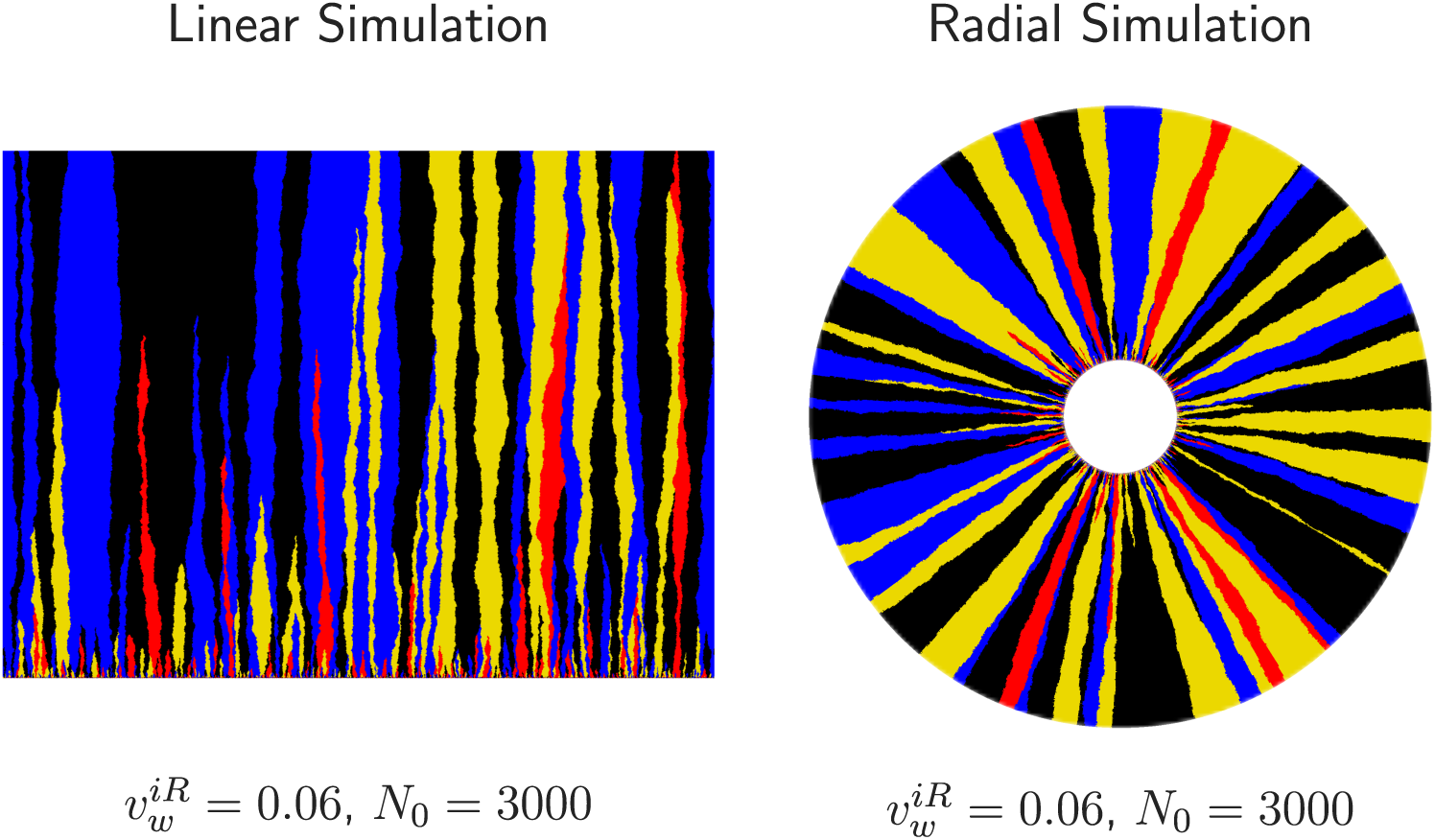
Simulations of linear (left) and inflating (right) range expansions grown for approximately 2000 generations. The black, yellow, and blue strains all sweep through the red strain with a wall velocity of 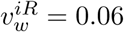. The initial concentration of all four strains was equal and *N*_0_ = 3000 was the number of cells that would fit on the initial front of both expansions. Note that the black, yellow, and blue sectors dominate over the red sectors at the end of these expansions.

### Calibrating the random walk model with experiment

To calibrate our experiments with the annihilating and coalescing random walk model, we measured *R*_0_, *D_w_*, and 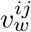; these parameters are illustrated in Fig. 1 and their ultimate values can be seen in Table 2. The values of these parameters were consistent between sets of plates made on different days.

#### Measuring *R*_0_

We defined the local fixation radius *R*_0_ as the radius where our image analysis package could correctly identify the inoculated fraction of multiple neutral strains, indicating that distinct sectors had formed and that our random walk model could describe their motion. For *R* < *R*_0_, our package predicted equal fractions of each strain due to the overlap of each channel in the homeland (see Figure 2). To determine *R*_0_, we inoculated radial expansions with three strains in unequal proportions; we used 10% of two strains and 80% of another. The minimum radius where we could accurately perceive fractions close to their initial values was *R*_0_ = 3.50 ± 0.05 mm as seen in supplementary Figure S2.

#### Measuring *D_w_*

Past work has found that *E. coli* colony domain walls fluctuate diffusively in certain conditions [18] and superdiffusively [8] in others. Creating a superdiffusive theory to describe the evolutionary dynamics of our system is beyond the scope of this paper. To obtain an effective diffusion constant *D_w_* and to test if the diffusive approximation adequately described our experimental dynamics, we fit the neutral Voter model’s prediction of heterozygosity (see Appendix S1), given by

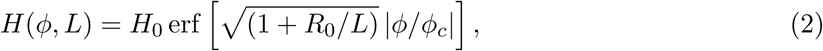

to experimental measurements. *H*_0_ is the heterozygosity for all when *L* = 0 for all *ϕ* (for *q* colors inoculated in equal fractions, *H*_0_ = 1 − 1/*q*) and 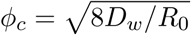 is a characteristic angular correlation length. The heterozygosity is the probability that two points separated by an angle of *ϕ* at a length expanded of *L* = *R* − *R*_0_ are occupied by different strains; it is a measure of spatial genetic diversity.

We fit *H*(*ϕ*, *L*) to our experimentally measured heterozygosity of two neutral strains (eCFP and eYFP) in three independent sets each with 14 range expansions. We averaged the heterozygosity over different experiments for each *L* as can be seen in Figure 5 (error bars were omitted for readability; the same figure with error bars can be found as Supplementary Figure S3). As we had previously measured *R*_0_ = 3.50 ± 0.05 mm, and *H*_0_ = 1/2 for two neutral strains inoculated at equal fractions, *D_w_* is the single free parameter in eq. (2). We consequently fit *D_w_* at each *L* with non-linear least-squares and found *D_w_* = (100 ± 5) × 10^−3^ mm after error propagation. The value of the diffusion constant is on the same order of magnitude as that from previous work [18].

**Fig 5.**
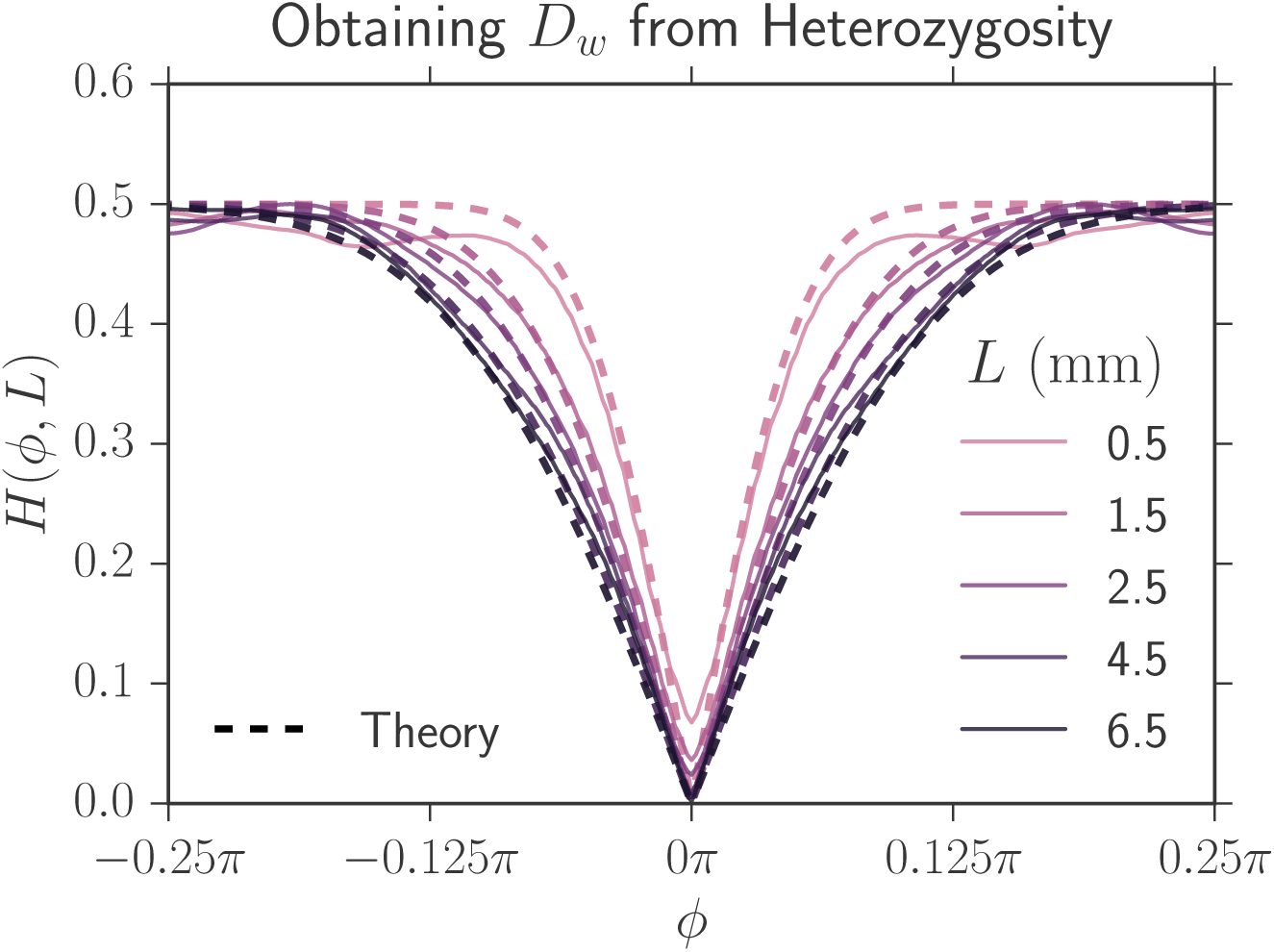
The heterozygosity correlation function *H*(*ϕ*, *L*) (solid lines) obtained by averaging the results of 14 neutral eCFP and eYFP expansions from one set of experiments at a variety of expansion distances *L* = *R* − *R*_0_. The dashed lines are the theoretical fits of the heterozygosity with a constant *D_w_* = (100 ± 5) × 10^−3^ mm. The theoretical curves track our experimental data, suggesting that a diffusive approximation to domain boundary motion is justified.

Figure 5 shows the Voter model’s fit (dashed lines) of the experimental heterozygosity (solid lines) using our values of *D_w_* and *R*_0_. The fit closely matches the experimental heterozygosity suggesting that a diffusive description of *E. coli* domain motion is justified. We use this value of *D_w_* for all strains. Unlike the Voter model and our simulations, the experimental heterozygosity at zero separation *H*(*L*, *ϕ* = 0) fails to vanish due to overlap between strains at domain boundaries; this effect is less pronounced at large radii because the effective angular width of boundaries decreased. The discrepancy between the theoretical and experimental heterozygosity is larger at small lengths expanded because the overlap between strains is larger; our image analysis is consequently less accurate.

#### Measuring 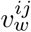

We used image analysis to directly quantify 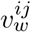 from the angular growth of more-fit sectors. Characteristic single sectors of each strain sweeping through the mCherry strain can be seen on the left side of Figure 6. In radial expansions, more fit strains should, on average, sweep logarithmic spirals through less fit strains at large lengths expanded, as verified in yeast expansions [17]. It can be shown that the average angular width of a sector of strain *i* sweeping through strain *j* is given by (see Appendix S1 for more details)

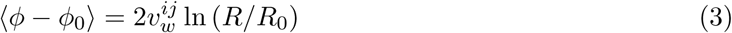

where *ϕ* is the angular width at radius *R* and *ϕ*_0_ is the initial angular width of the domain at *R*_0_. 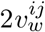 can thus be extracted from the slope of a linear regression fit of 〈*ϕ* − *ϕ*_0_〉 vs. ln(*R*/*R*_0_) as seen on the right side of Figure 6.

**Fig 6.**
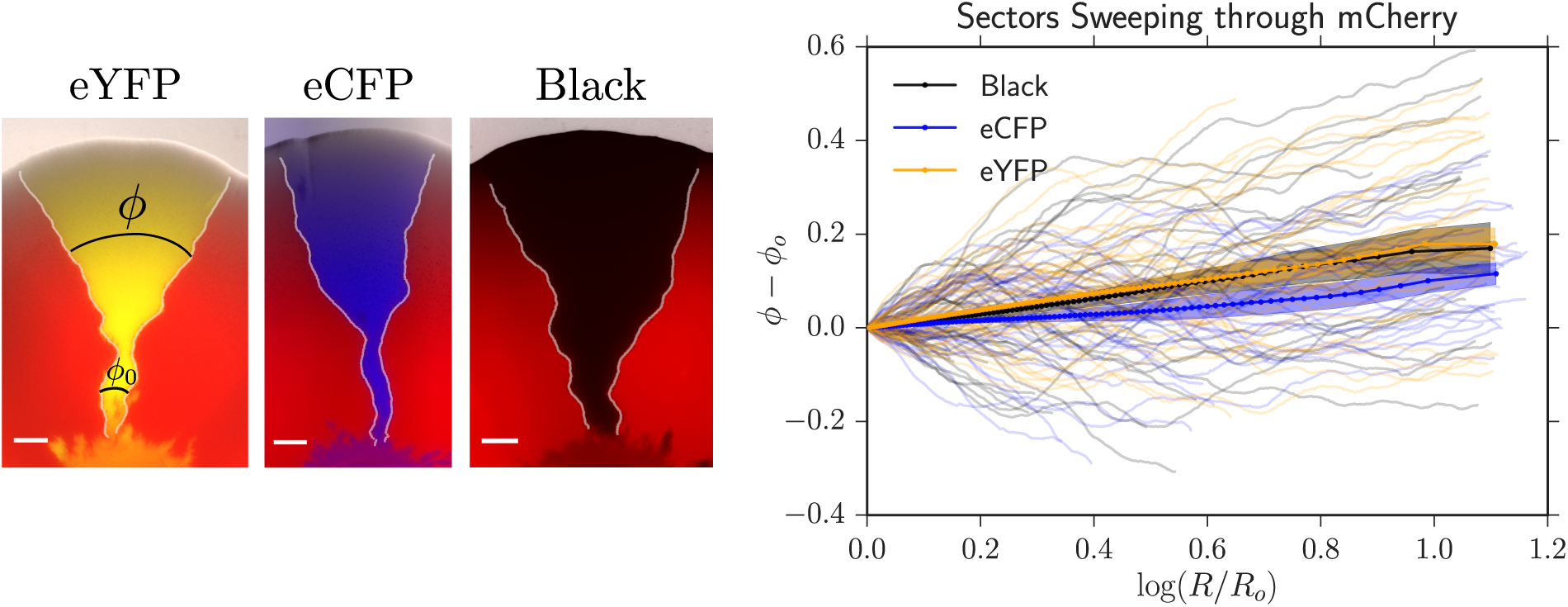
*Left:* Single sectors of our eYFP, eCFP, and black strains sweeping through the less fit strain mCherry. The scale bar is 1 mm. The white lines are the positions of the domain walls located with our image analysis package [34]. We tracked the angular growth of sectors sweeping through a less fit strain, *ϕ* − *ϕ*_0_, as a function of ln (*R*/*R*_0_) to obtain 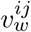. *Right:* 40 traces of each strain sweeping through mCherry. The translucent lines are the individual traces, the solid lines are the mean angular growth 〈*ϕ* − *ϕ*_0_〉, and the shaded area is the standard error of the mean. The slope of the mean angular growth is 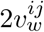.

By tracking domain walls directly, we found that more fit strains swept through the less fit mCherry strain with a wall velocity of 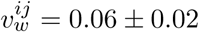. The wall velocity was significantly smaller than expected from the basal independent expansion velocities of our strains (Table 1); potential explanations for this phenomenon are discussed in Appendix S2. The magnitude of the velocities were consistent between experiments (using 40 single sectors on three sets of plates) but were too imprecise to be predictive in our models. We consequently developed a more precise technique to measure the velocities 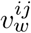 using a comparison to simulation in the main text.

## Results and discussion

In this paper, we experimentally quantify and theoretically predict the average fraction, the two-point correlation functions, and the relative rate of annihilations and coalescences per domain wall collision for four competing *E. coli* strains with different expansion velocities. The different strains of *E. coli* represent competing alleles. In this section, we first frame our results by introducing key parameters governing the evolutionary dynamics. We then report our experimental measurements and show how our annihilating and coalescing random-walk model can utilize the key parameters to predict the dynamics of the four competing strains.

### Key parameters

What key combinations of parameters from our annihilating and coalescing random walk model govern the evolutionary dynamics? Our goal is to describe the dynamics as a function of length expanded by our colonies *L* = *R* − *R*_0_ with *R*_0_ the initial radius where domain walls form, the domain wall diffusion coefficient per length expanded *D_w_* (units of length), and all wall velocities per length expanded 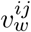 (dimensionless). The two-point correlation functions must include an additional independent variable: the angular distance *ϕ* between strains.

Investigating the width of a single sector of a more fit allele sweeping through a less fit allele, as illustrated on the right of Figure 1, reveals important parameter combinations (see Appendix S1 for additional details). In a linear expansion, the deterministic, selection-induced growth of a sector of genotype *i* sweeping through a less fit genotype *j* will scale as 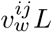 while its diffusive growth will scale as 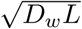. At short lengths expanded, diffusion will thus dominate deterministic growth, and at larger lengths selection will dominate diffusion. A crossover expansion length 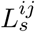 [2,9,32] beyond which selection dominates follows by equating the deterministic and diffusive growth,

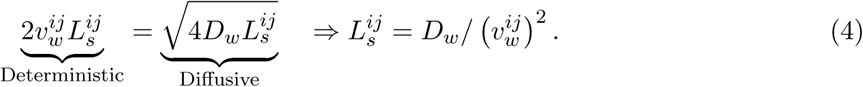

The factor of 2 in front of 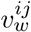 and 4 in front of *D_w_* arises because we are monitoring the distance between *two* domain walls (i.e. a sector); similar arguments can be applied to describe the motion of individual walls. 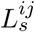 is the characteristic length that the colony must expand in order for selection to dominate over diffusion for strain *i* sweeping through strain *j* and acts as the first key parameter.

Upon repeating this argument for domains on a radially inflating ring (see Appendix S1), we identify 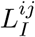 [32, 42] as the inflationary analog of 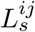: the expansion length beyond which selection dominates over diffusion, and find

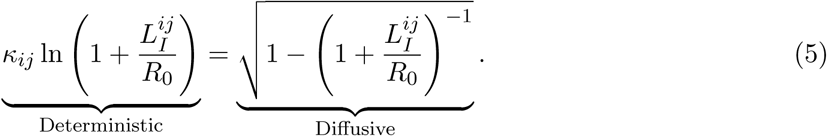

*κ_ij_* is a dimensionless prefactor that can be thought of as an “inflationary selective advantage” controlling the expansion length at which selection dominates over diffusion and is given by

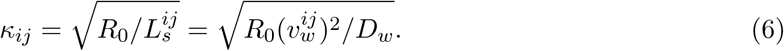

Fig. 7 illustrates the importance of *κ_ij_*; it displays the ratio of the inflationary to the linear selection length scale 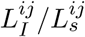 as a function of *κ_ij_* from the numerical solution of eq. (5). We find that the ratio of the length scales has the asymptotic behavior

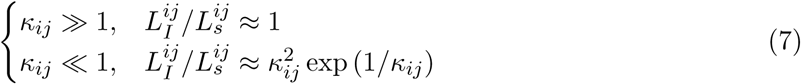

Thus, if *κ_ij_* ≫ 1, inflation can be ignored, as the inflating selection length scale approaches the linear selection length scale. In contrast, if *κ_ij_* ≪ 1, the inflationary selection length will be many times larger than the linear selection length scale [32]. Therefore, as *κ_ij_* becomes smaller, diffusion dominates over selection for a larger length expanded. *κ_ij_* is the second key parameter describing the dynamics of our system.

**Fig 7.**
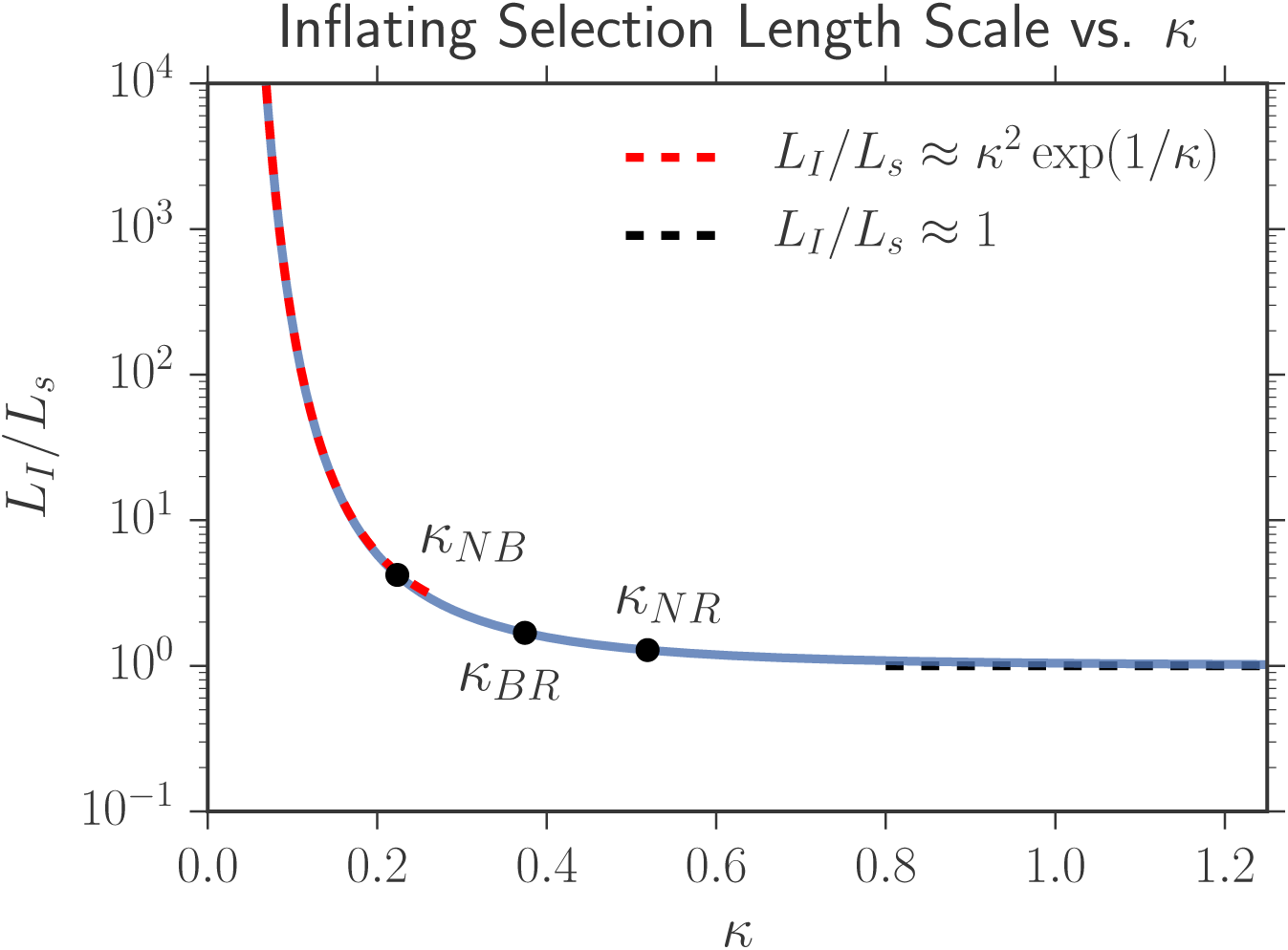
Numerical solution of eq. (5) describing how *L_I_* varies as a function of *κ*; asymptotic approximations are overlaid. If *κ* ≳ 1, inflation does not appreciably slow selective sweeps as *L_I_* approaches the linear selection length scale *L_s_*. In contrast, if *κ* ≪ 1, the inflationary selection length scale *L_I_* will be many times larger than the linear selection length scale *L_s_*, indicating that diffusion will dominate for a much larger length expanded. The three black points correspond to measurements of the *κ_ij_* that govern the dynamics of our competing strains (see Table 3 and the surrounding discussion); *N* stands for the two selectively neutral strains (eCFP and eYFP), *B* for black, and *R* for mCherry (red). Inflation slows the sweep of the neutral strains through black the most because *κ_NB_* is smaller than *κ_BR_* and *κ_NR_*; smaller values of *κ* have a larger value of *L_I_*/*L_s_* indicating that it takes longer for deterministic motion to dominate over diffusive motion.

The third and final key parameter is the characteristic angular correlation length 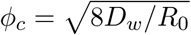 between selectively neutral genotypes and appeared above in eq. (2). This parameter arises naturally when analytically calculating the neutral two-point correlation functions from the Voter model (see eq. 10). The parameter also has an intuitive description. When moving into polar coordinates, the angular diffusion coefficient *D_ϕ_* is related to the standard linear domain wall diffusion coefficient by *D_ϕ_* = *D_w_*/*R*. The characteristic scale for the radius is *R*_0_ and the average diffusive growth of domains should scale as 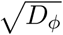; it thus makes sense that 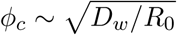.

We have now identified the three key parameters that govern the evolutionary dynamics of our colonies. 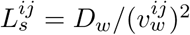 is the length that a linear expansion must grow in order for selection to dominate over diffusion for strain *i* sweeping through strain *j*, 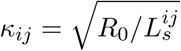 controls how quickly selection dominates over diffusion in radially inflating expansions, and 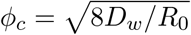 sets the characteristic angular correlation length between selectively neutral genotypes. These key parameters are listed in Table 2.

**Table 2.** Parameters used in the annihilating and coalescing random-walk model. We experimentally quantified *R*_0_, *D_w_*, and *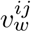* so that we could compare experiments to the model.

| Parameter | Value | Description |
| --- | --- | --- |
| Experimentally Measured |  |  |
| $R_0$ | $3.50 \pm 0.05$ mm | Initial radius where domain walls form |
| $D_w$ | $(100 \pm 5) \times 10^{-3}$ mm | Diffusion constant per length expanded |
| $v_w^{ij}$ | $0.06 \pm 0.02$ | Wall velocity: transverse displacement per length expanded.<br>Only observable for strains sweeping through mCherry. |
| Key Parameters |  |  |
| $L_s^{ij}$ | $D_w/(v_w^{ij})^2$ | Length expanded when selection dominates diffusion for linear expansions |
| $\kappa_{ij}$ | $\sqrt{R_0/L_s^{ij}}$ | Controls how quickly selection dominates diffusion in radial expansions |
| $\phi_c$ | $\sqrt{8D_w/R_0}$ | Characteristic angular correlation length between strains |
| Independent Variables |  |  |
| $R$ | – | Radius of the colony |
| $L$ | $R - R_0$ | Colony length expanded |
| $\phi$ | – | Polar coordinate angle |

### Experimental measurements

We now introduce our experimental measurements of the average fraction of each strain, the two-point correlation functions between strains, and the relative rates of annihilations and coalescences as a function of length expanded. In this section, we use neutral theory as a null model; deviations from these predictions illustrate the impact of selection.

#### Average fractions in three-color expansions

The average fraction of strain *i* at a length expanded of *L* = *R* − *R*_0_ is given by

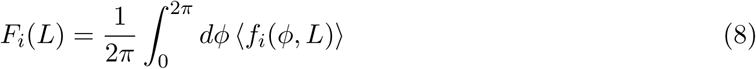

where *f_i_*(*ϕ*, *L*) is the local fraction of strain *i* at angle *ϕ* and length *L* (i.e. at a pixel specified in polar coordinates by *ϕ* and *L*). The angular brackets represent an average over many range expansions and *f_i_* is normalized such that 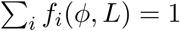 as discussed in the Image Analysis section. In the neutral case, the average fraction of each strain should equal their inoculated fractions and should be independent of length expanded. Selection forces the average fractions of less fit strains to decrease.

We measured the average fraction versus radial length expanded in two separate sets of experiments where we inoculated different fractions of our eYFP, eCFP, and mCherry strains. In one experiment, we inoculated the eYFP, eCFP, and mCherry strains with equal initial fractions of 33% while the other we inoculated 80% of the mCherry strain and 10% each of the eCFP and eYFP strains. We conducted 20 replicates in each case and calculated the average fraction of each strain using our image analysis package. Figure 8 displays the trajectories of the 20 expansions and the mean trajectory (the average fraction) as ternary composition diagrams for both sets of initial conditions [43].

**Fig 8.**
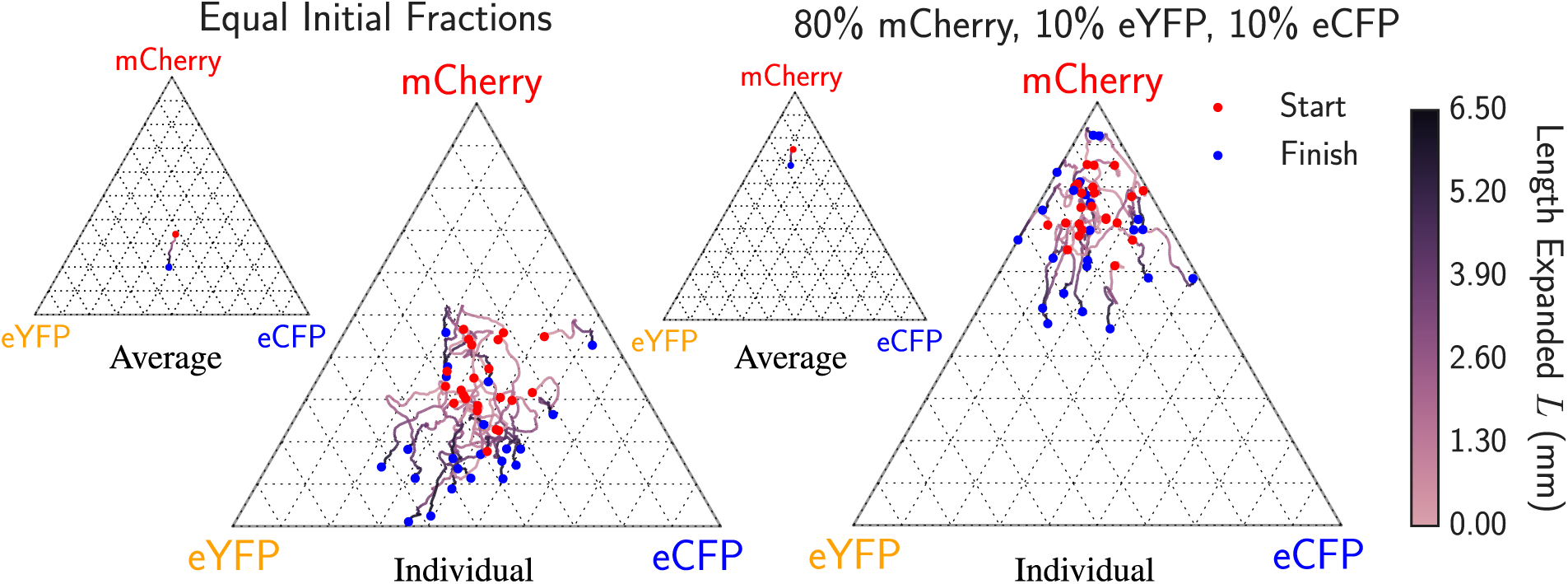
Average fraction of each genotype as a function of length expanded for 20 radial expansions each when equal fractions of eCFP, eYFP, and mCherry were inoculated (left) and when 10% eCFP, 10% eYFP, and 80% mCherry were inoculated (right). The red dot indicates the composition at the radius *R*_0_ = 3.50 mm where distinct domain walls form and the blue dot indicates the composition at the end of the experiment. The red dots are dispersed about the initial inoculated fractions due to the stochastic dynamics at the early stages of the range expansions when *R* < *R*_0_. The highly stochastic trajectories illustrate the importance of genetic drift at the frontier in the *E. coli* range expansions. The smaller ternary diagrams display the average fraction over all expansions vs. length expanded for each set of experiments. For both initial conditions, we see a small systematic drift away from the mCherry vertex indicating that the mCherry strain has a lower fitness, qualitatively agreeing with the selective advantages of Table 1. Note that two replicates on the right resulted in the complete extinction of eCFP due to strong spatial diffusion, indicated by the trajectories pinned on the absorbing line connecting the eYFP and mCherry vertices.

In both sets of experiments, we observed a systematic drift away from the mCherry vertex as a function of radius as illustrated by the mean trajectories shown as insets. We witnessed two cases where the 10% initial inoculant of the eCFP strain became extinct, represented by the pinning of trajectories to the absorbing boundary connecting the eYFP and mCherry vertex, a consequence of the strong genetic drift at the frontiers of our *E. coli* range expansions. These measurements indicate that the mCherry strain was less fit than the eCFP and eYFP strains, consistent with the order of the radial expansion velocities.

#### Two-point correlation functions

We measured the two-point correlation functions given by

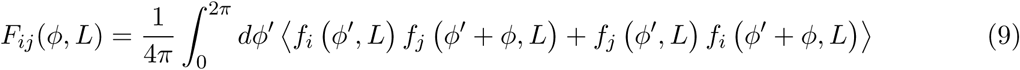

where *f_i_*(*ϕ*, *L*) is again the local fraction of strain *i* at angle *ϕ* and expansion length *L*. *F_ij_* gives us the probability that strain *i* is located at an angular distance of *ϕ* away from strain *j* at a length expanded *L*. Note that *F_ij_* = *F_ji_* and *F_ij_*(*ϕ*) = *F_ij_* (−*ϕ*). Although the average fraction is constant in the neutral case, the two-point correlation functions broaden due to the coarsening of genetic domains [2]. Neutral *q*-color Voter models analytically predict the form of the two-point correlation functions [2] as discussed in Appendix S1 as

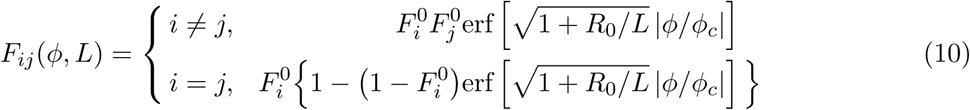

where 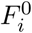 is the initial fraction inoculated of strain *i* and 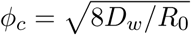 is the characteristic angular correlation length discussed above. If we evaluate *ϕ_c_* using our experimental values of *D_w_* and *R*_0_, we find that *ϕ_c_* = 0.48 ± 0.01 radians; *ϕ_c_* is the same as that from the heterozygosity calculation in eq. (2). As the colony expands, *R*_0_/*L* → 0 and the two-point correlation functions will “freeze” with a characteristic angular width of *ϕ_c_*.

Deviations from neutral predictions are caused by selection and can be understood in the limit of both large and small angular separations. For large angular separations relative to the characteristic angular width, two-point correlation functions will factorize and plateau at the value *F_ij_* ≈ *F_i_F_j_* provided that *ϕ_c_* ≪ 2*π*. Selection can be identified by comparing the experimentally measured plateau value to the neutral prediction value of 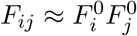, the product of the initial fractions inoculated of strains *i* and *j*. Furthermore, in the limit of zero angular separation, it is known that *∂_ϕ_F_ij_* measures the density of *ij* domain walls [2] (note that *i* ≠ *j*). In general, if strain *i* is less fit than the other strains, it will have fewer domain walls, decreasing the domain-wall density and thus the slope near *ϕ* = 0.

We measured the correlation functions between each pair of strains in three sets of experiments where we inoculated equal well-mixed fractions of the eCFP, eYFP, and black strains, then eCFP, eYFP, and mCherry, and then finally all four strains. We conducted 20 replicates of each experiment, measured all two-point correlation functions at the final radius of *R* = 10 mm corresponding to a length expanded of *L* = *R* − *R*_0_ = 6.5 mm, and averaged the results. In Figure 9, we plotted the neutral correlation function prediction (10) as a control, and compared it to the experimentally measured correlation functions.

**Fig 9.**
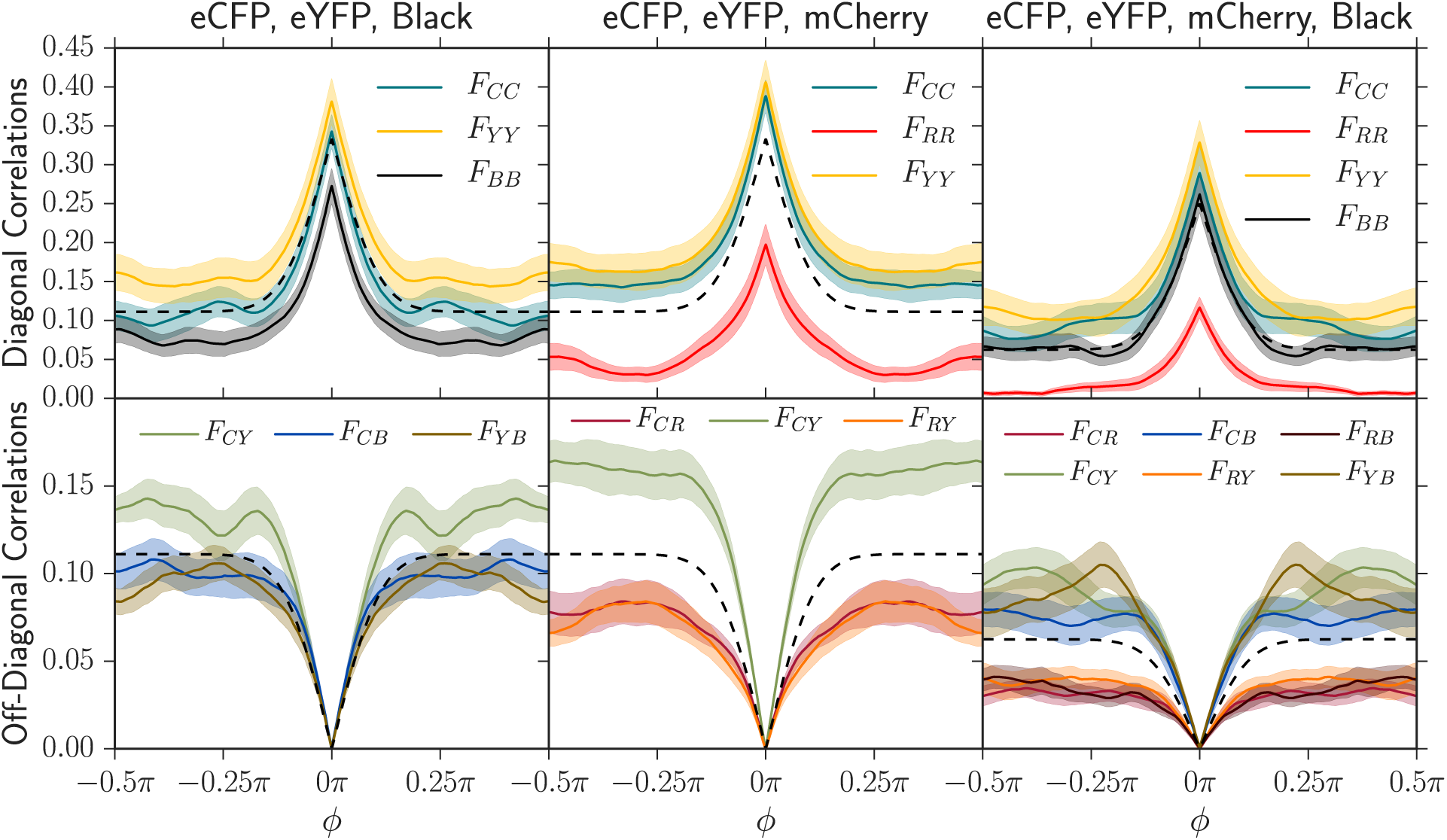
Two-point angular correlation functions *F_ij_* (*L*, *ϕ*) at a length expanded of *L* = *R* − *R*_0_ = 6.5 mm (*R*_0_ = 3.5 mm) in three sets of experiments where we inoculated 20 replicates with equal fractions of our eCFP, eYFP, and black strains (left), then eCFP, eYFP, and mCherry (center), and finally all four strains (right). The shaded regions in these plots indicate standard errors of the mean. Using the measured diffusion coefficient *D_w_* and initial radius where domain walls form *R*_0_, we also plot the theoretical two-point correlation functions from the neutral Voter model (black dashed line) from eq. 10. The colors of each plotted correlation function were chosen to correspond to their composite strain colors; for example, two-point correlation correlation functions associated with mCherry were red or were blended with red for the panels on the right. The subscripts correspond to the color of each strain: *C* = eCFP, *Y* = eYFP, *R* = mCherry, and *B* = Black. As judged by the magnitude of the deviation from neutral predictions, the black strain has a small selective disadvantage relative to eCFP and eYFP and that the mCherry strain has an even greater disadvantage, in agreement with the independent radial expansion velocities in Table 1.

The two-point correlation functions in the experiment between eCFP, eYFP, and and the black strains (first column of Figure 9) are consistent with the order of radial expansion velocities (fitnesses) in Table 1. The correlation between the eCFP and eYFP strains plateaued at a higher value than the correlation between eCFP and black and the eYFP and black, indicating that the eCFP and eYFP strains were indeed more fit. The self-correlation for the black strain, *F_BB_*, plateaued at the smallest value relative to eCFP and eYFP.

In contrast, combining eCFP, eYFP, and mCherry in one set of experiments and all four strains in another revealed that mCherry had a larger fitness defect. Correlation functions including mCherry always plateaued at a significantly smaller value than correlation functions excluding it. Furthermore, correlation functions involving the mCherry strain had a smaller slope at zero angular separation, indicating that less mCherry domain walls were present and that the mCherry strain was less fit than the others. The two-point correlation functions were thus consistent with the black strain having a small selective disadvantage relative to eCFP and eYFP and the mCherry strain having a larger disadvantage relative to all others.

#### Annihilation asymmetry

The last quantity we measured was the relative rate of annihilations and coalescences per domain wall collision; examples of annihilations and coalescences can be seen in Figure 1. Although little is known about the relative rates of annihilations and coalescences in the presence of selection, analytical results are available for the neutral case (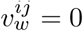). If *q* (an integer) neutral alleles are inoculated at random locations with *equal* initial proportions on a one-dimensional lattice, the probability of obtaining an annihilation per domain wall collision is given by

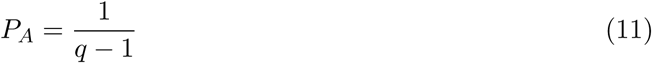

and the probability of obtaining a coalescence per collision is given by [27–29]

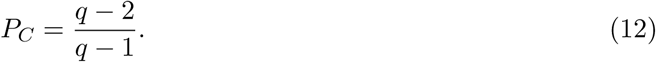

One can easily derive these formulas; given that strain *i* is to the left of two colliding domain walls and strain *j* is between them, we can ask: “what is the probability that strain *k* to the right of the walls is the same as strain *i* (annihilation) or is *not* strain *i* or *j* (coalescence)?” Note that although the rate of annihilations and coalescences decreases with time due to coarsening, the probabilities per collision *P_A,C_* are independent of the length expanded *L*. In the presence of selection, however, the average global fraction of each strain will change with length expanded, making *P_A_* and *P_C_ L*-dependent.

To succinctly quantify the difference between the annihilation and coalescence probabilities, we define the “annihilation asymmetry” Δ*P*(*L*) = *P_A_*(*L*) − *P_C_*(*L*) as the difference in probability of obtaining an annihilation versus a coalescence per collision at a distance expanded of *L*. If *q* neutral colors are inoculated in equal fractions, we find, using equations (11) and (12),

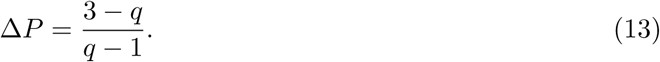

Thus, for the case of *q* neutral colors in equal proportions, we have lim*_q_*_→∞_ Δ*P*(*q*) = −1 (only coalescences), Δ*P*(*q* = 3) = 0 (equal numbers of annihilations and coalescences), and Δ*P*(*q* = 2) = 1 (only annihilations). The quantity Δ*P* thus provides a simple way to characterize the annihilation/coalescence difference in a single curve that varies smoothly between −1 and 1 as 2 ≤ *q* < ∞. In Appendix S1 we develop and discuss the case of non-equal proportions (supplementary equations (S1.5)–(S1.7)). For this case it is then useful to define a “fractional q” by inverting equation (13) to read *q* = (3 + Δ*P*)/(1 + Δ*P*).

To quantify the annihilation asymmetry, we examined the average cumulative difference in annihilations and coalescences vs. the average cumulative number of domain wall collisions as colonies expanded; Δ*P* is given by the slope of this quantity and can be seen in Figure 10 (see Supplementary Figure S4 for a display of cumulative count vs. length expanded). Regardless of which strains were inoculated and their selective differences, our results were consistent with the neutral theory prediction in equation (13) for *q* = 2, *q* = 3, and *q* = 4 as judged by the overlap of the black dashed line with the shaded standard error of the mean in each case. Δ*P* appeared to be constant as a function of length. We also tested an initial condition where we inoculated strains in unequal proportions: we inoculated 10% of eCFP and eYFP and 80% of mCherry. This experiment again matched the neutral prediction of Δ*P* ≈ 0.51 (and correspondingly *q* ≈ 2.33) within error. We discuss why certain observables, like the average fraction and two-point correlation functions, showed signatures of selection while others, like the annihilation asymmetry, did not in the Predicting Experimental Dynamics section.

**Fig 10.**
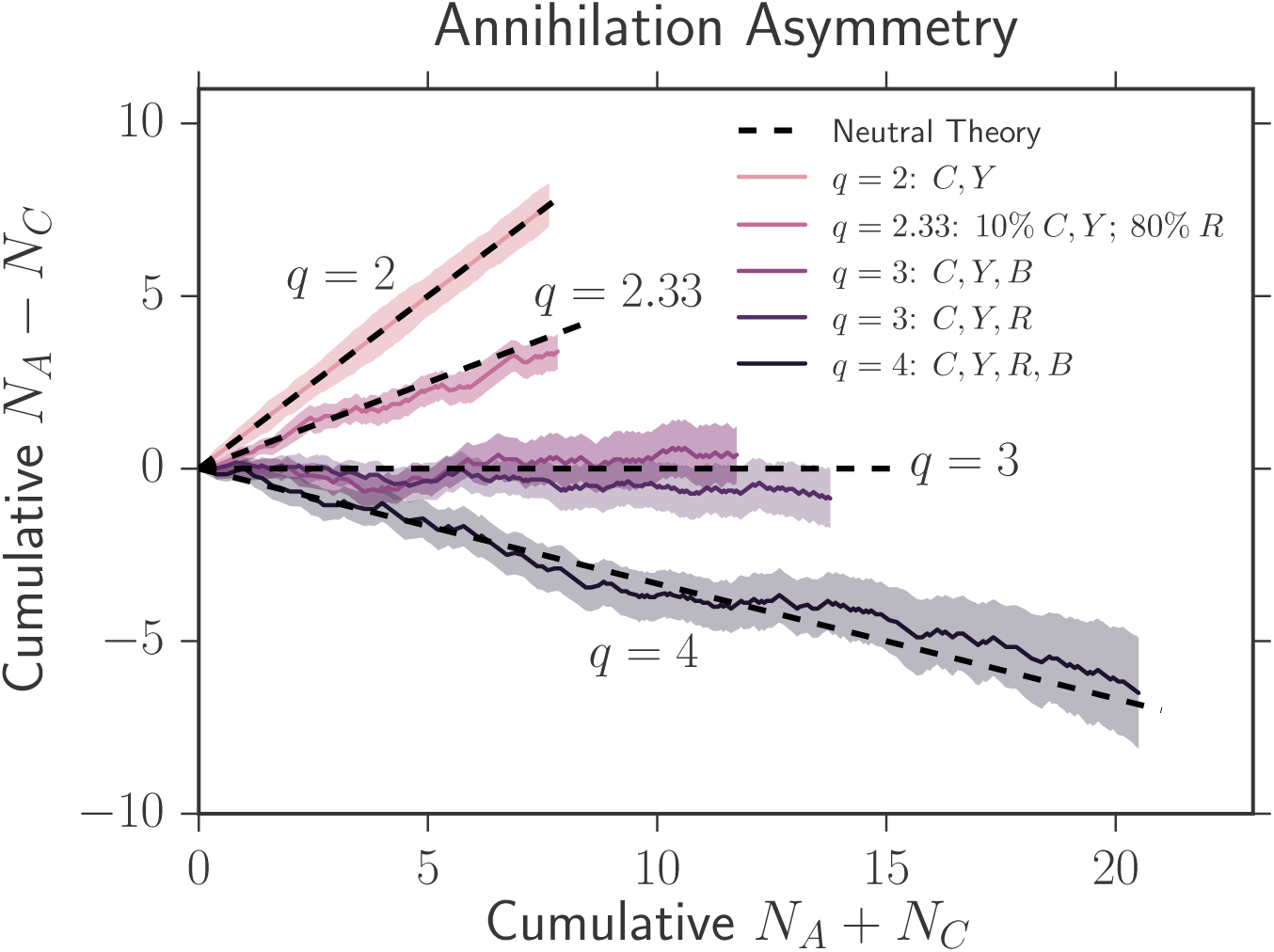
Average cumulative difference in annihilations and coalescences vs. the average cumulative number of domain wall collisions as colonies expand. The slope of this plot gives the annihilation asymmetry Δ*P*. The shaded regions represent the standard error of the mean between many experiments. We use the notation *C* = eCFP, *Y* = eYFP, *B* = black, and *R* = mCherry. Despite the presence of selection, Δ*P* was consistent with the standard neutral theory prediction of eq. (13) Δ*P* = (3 − *q*)/(*q* − 1) for *q* = 2, *q* = 3, and *q* = 4 (equal initial fractions of *q* strains), as judged by the overlap of the black dashed lines with the shaded areas in every case. We also explored an initial condition where we inoculated unequal fractions of three strains; we inoculated 10% of eCFP and eYFP and 80% of mCherry. Our experiments agreed with the prediction of Δ*P* ≈ 0.51, or an effective *q* ≈ 2.33, from the neutral theory developed in supplementary equations (S1.5)–(S1.7). We discus why the annihilation asymmetry did not show signatures of selection in the Predicting Experimental Dynamics section.

### Collapsing the evolutionary dynamics with simulation

We used our simulations to investigate the effect of the model’s input parameters *R*_0_, *D_w_*, and the set of all 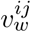 on our three observables: the average fraction of each strain, the two-point correlation functions between strains, and the relative rate of annihilations and coalescences per domain wall collision. We simulated both linear and radial expansions using the algorithm discussed in the Simulation Section which has a wall diffusion constant per length expanded of *D_w_* = *a*/2, where *a* is the cell diameter, and a wall velocity per length expanded of 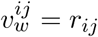, where *r_ij_* is the probabilistic bias that a domain wall has of jumping in a certain direction. For the radial expansions the initial radius was given by *R*_0_ = *N*_0_*a*/(2*π*), where *N*_0_ is the initial number of individuals at the population perimeter when domain walls formed, and the radius was inflated appropriately. In linear simulations, radius was held fixed, so the simulation was effectively carried out on a cylinder [2]. In every simulation, we kept *D_w_*, *a*, and the expansion velocity constant as in our experiments. We varied 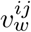 and *R*_0_ by altering the simulation parameters *r_ij_* and *N*_0_ respectively.

The three key parameters governing the evolutionary dynamics of our system (discussed in the Key Parameters section) are 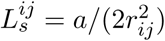, the length that a linear expansion must grow in order for selection to dominate over diffusion for strain *i* sweeping through strain *j*, 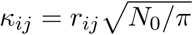, which controls how much selection dominates over diffusion in radially inflating expansions, and 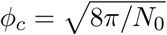, the characteristic angular correlation length between strains.

We first simulated *q* = 3 competing strains where two neutral strains swept through a third less fit strain with wall velocity *v_w_*, similar to our experiments with two neutral colors (eCFP and eYFP) and the less fit mCherry strain. The three strains were inoculated in equal proportions. Note that in this simulation, there was only one non-zero *v_w_* and consequently one 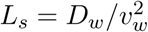 and one 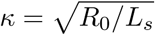. We varied *v_w_* from 10^−3^ ≤ *v_w_* ≤ 10^−1^ and *N*_0_ from 10^2^ ≤ *N*_0_ ≤ 10^5^ and computed the average fraction *F* of the less fit strain and the annihilation asymmetry Δ*P*. We found that both *F* and Δ*P* from simulations with identical *κ*, despite different values of *R*_0_ and *v_w_*, collapsed if the length traveled *L* was rescaled by *L_s_* as seen in Figure 11. Each curve in Figure 11 consists of 6 collapsed simulations with unique values of *R*_0_ and *v_w_* but with the same value of *κ*. Further simulations revealed that the the two-point correlation functions *F_ij_* could be collapsed from simulations with identical *κ* if *L* was rescaled by *L_s_* and *ϕ* was rescaled by 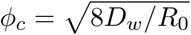 provided *ϕ_c_* ≪ 2*π* (see Supplementary Figure S7).

**Fig 11.**
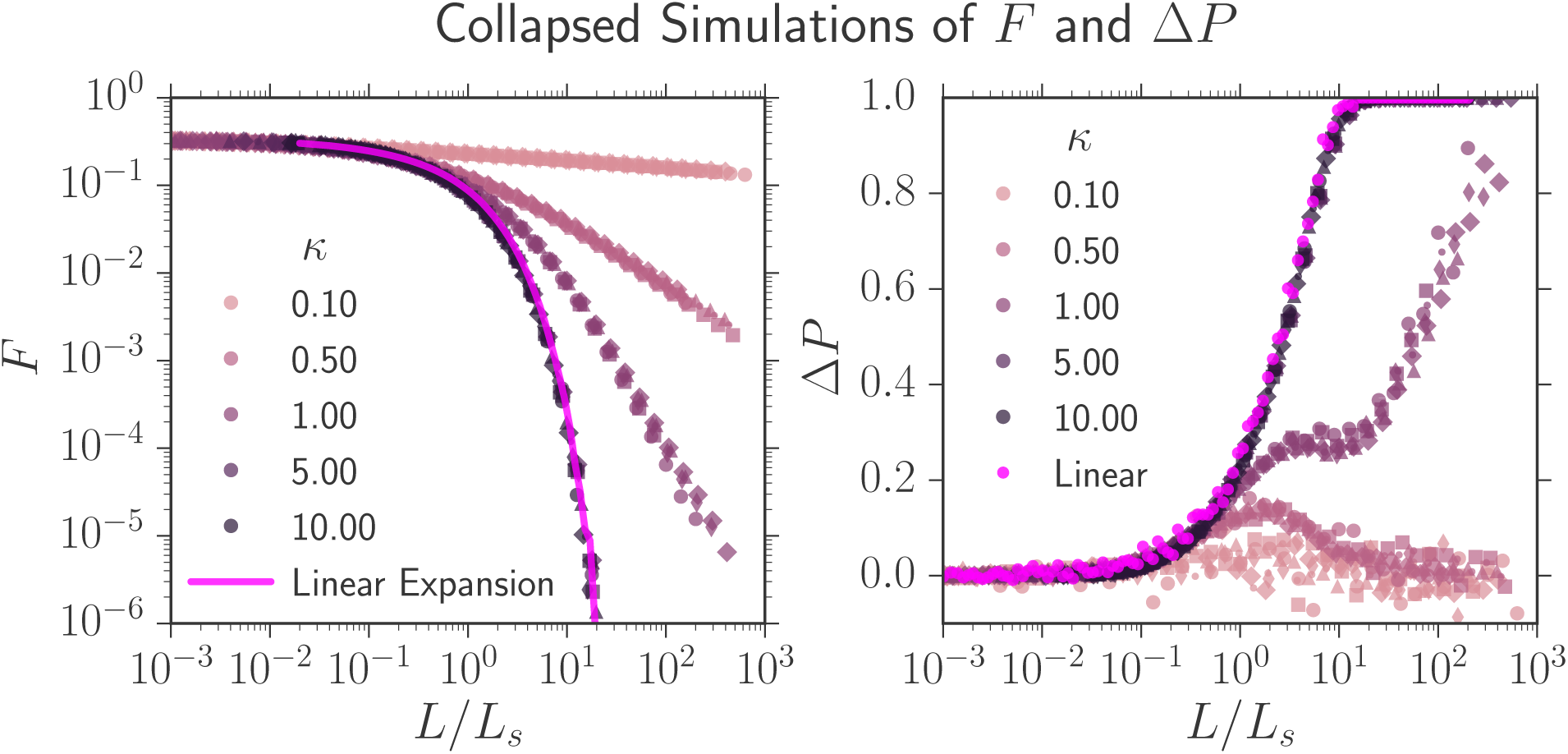
Simulations of the average fraction *F* of a less fit strain and the annihilation asymmetry Δ*P* collapsed onto universal curves parameterized by 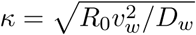.Two neutral strains swept through a less fit strain (its fraction given by *F*) with a wall velocity *v_w_* ; each strain was inoculated in equal proportions and the colony’s initial radius was *R*_0_. If simulations had identical *κ*, despite different values of *R*_0_ and *v_w_*, both *F* and Δ*P* could be collapsed if the length traveled *L* was rescaled by 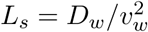,the linear selection length scale.Each universal curve at a fixed *κ* consists of 6 simulations with different values of *v_w_* and *R*_0_; each set of parameters has a different marker. As *κ* decreased, inflation slowed the selective sweep of the more fit strains through the less fit strain as illustrated by the slower decrease of *F*. Δ*P* transitioned from 0 to 1 as the number of strains present in the expansion decreased from *q* = 3 to *q* = 2 (the less fit strain was squeezed out); this is expected from eq. (13), Δ*P* = (3 − *q*)/(*q* − 1).

We now consider the collapsed curves *F*(*L*/*L_s_*, *κ*) and Δ*P*(*L*/*L_s_*, *κ*) as a function of the parameter *κ* as seen in Figure 11. *κ* had a pronounced effect on both quantities. For 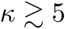 the dynamics of *F* and Δ*P* approached the dynamics of a linear expansion at all *L*/*L_s_*, illustrated by the bright pink line on the left and the bright pink dots on the right of Figure 11; the more fit strain swept so quickly through the less fit strain that the colony’s radial expansion could be ignored. As *κ* decreased, the less fit strain was squeezed out more slowly due to the inflation of the frontier, resulting in slower transitions from *q* = 3 to *q* = 2 colors and consequently slower transitions from Δ*P* = 0 to Δ*P* = 1. For *κ* ≪ 1, Δ*P* barely shifted from 0 over the course of the simulation. Interestingly, Δ*P* peaked at a finite *L*/*L_s_* for small *κ*; it is not clear what causes this effect, but it may be related to the transition from linear to inflation-dominated dynamics as *L* increases.

Additional simulations revealed that for expansions composed of many strains with different fitnesses (multiple 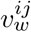) and consequently various *κ_ij_*, all of our observables (*F*, Δ*P*, and *F_ij_*) could again be collapsed onto a master curve by rescaling *L* by any one of the selection length scales (i.e. 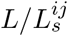) and by rescaling *ϕ* by *ϕ_c_*; the set of *κ_ij_* specified the master curve. An example of a simulation that exhibits collapsed dynamics for a wide range of parameters can be seen in Figure S8 in the Supplementary Material.

To summarize the results of this section, we found that we could collapse the average fraction *F*, annihilation asymmetry Δ*P*, and the two-point correlation functions *F_ij_* by

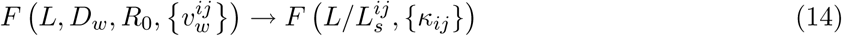

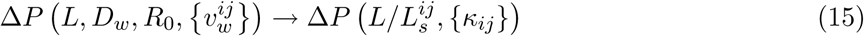

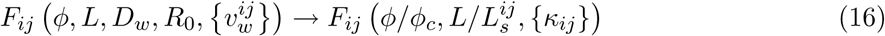

where the brackets indicate a *set* of variables parameterized by *i* and *j* (i.e. {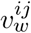} represents the set of all *ij* wall velocities). As long as *L* was rescaled by any selection length scale 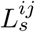 and *ϕ* was rescaled by the characteristic angular correlation length *ϕ_c_*, the set of {*κ_ij_*} completely dictated the evolutionary dynamics.

### Predicting experimental dynamics via collapsed simulations

A major goal of this paper is to test if the annihilating and coalescing random-walk model can predict the experimental evolutionary dynamics of our four competing strains (alleles) with different fitnesses (radial expansion velocities). To the best of our knowledge, analytical results from the random-walk model are unavailable; we consequently used our simulations to predict the dynamics. We needed the experimental measurements of *D_w_*, *R*_0_, and 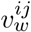 in order to set the simulation parameters *a*, *d*, *r_ij_*, and *N*_0_. Uniquely fitting four simulation parameters is challenging, however, and noisy; as the number of free parameters increases, simulation results become hard to interpret.

We consequently took an alternative approach to calibrate our simulations to experiment. In the last section, we found that the simulation’s dynamics could be collapsed onto master curves for a fixed set 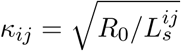 by rescaling the length expanded *L* by any 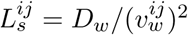 and by rescaling *ϕ* by 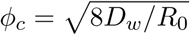. These simulated master curves were invariant to the alteration of simulation parameters provided that *κ_ij_* remained the same. Therefore, to avoid specifically setting simulation parameters, we *experimentally* evaluated 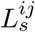, *ϕ_c_*, and *κ_ij_*, and tested if the rescaled (*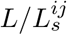* and *ϕ*/*ϕ_c_*) experimental dynamics agreed with the universal simulated curve at the same set of *κ_ij_*.

We had previously determined *R*_0_ = 3.50 ± 0.05 mm, *D_w_* = (100 ± 5) × 10^−3^ mm, and 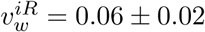 as seen in Table 2. In principle, these values should allow us to totally calibrate the three key parameters. For example, 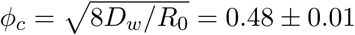. The value of 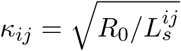 followed from the measurement of 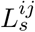 using the known value of *R*_0_. Unfortunately, the final parameter 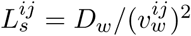 was more difficult to calibrate. Using 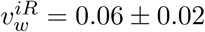, we found that 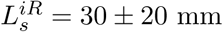; the error on this value was too large for it to be predictive in our simulations. Furthermore, we were unable to accurately measure the wall velocity of the eCFP/eYFP strains sweeping through the black strain, and consequently could not calculate the corresponding selection length scale.

To overcome these issues, we created a new, more precise technique to fit *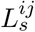* (and thus 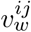) that compared universal simulated correlation functions to experimental correlation functions extracted from pairwise competitions between strains. After introducing this technique, we show that the resulting set of key parameters can be used to predict the evolutionary dynamics of the four strains when grown together in an independent experiment.

#### Fitting *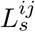* via collapsed two-point correlation functions

As our eCFP and eYFP strains were neutral within error, we treated our system as composed of one neutral (*N*) eCFP/eYFP strain, a red (*R*) mCherry strain, and a black (*B*) strain (*q* = 3 colors). As the eCFP/eYFP expanded faster than the black followed by the mCherry strain, we needed to quantify 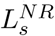, 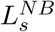, and 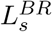.

To fit 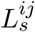 more precisely than that from our direct measurement of wall velocity, we competed pairwise combinations of strains in range expansions (i.e. the eCFP/eYFP strain vs. mCherry) and calculated the two-point correlation functions *F_ij_*(*L*, *ϕ*) at the maximum length expanded of *L* = 6.5 mm. As there were only two competing strains, there was only one *L_s_*. To find this value, we rescaled our experimental length expanded *L* by *L_s_* (the parameter we were fitting) and *ϕ* by *ϕ_c_* (previously calculated) and compared the rescaled experimental data to the simulated universal functions with the corresponding parameters of *L*/*L_s_*, *ϕ*/*ϕ_c_*, and 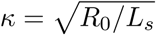. Figure 12 illustrates this procedure by displaying an experimentally rescaled two-point correlation function *F_NR_* (the solid red line) between our eCFP/eYFP strain (*N*) and our mCherry strain (*R*), which were inoculated at fractions of 2/3 and 1/3, respectively, at a length expanded of *L* = 6.5 mm, and simulated universal correlation functions corresponding to different values of *L_s_* (dashed lines).

**Fig 12.**
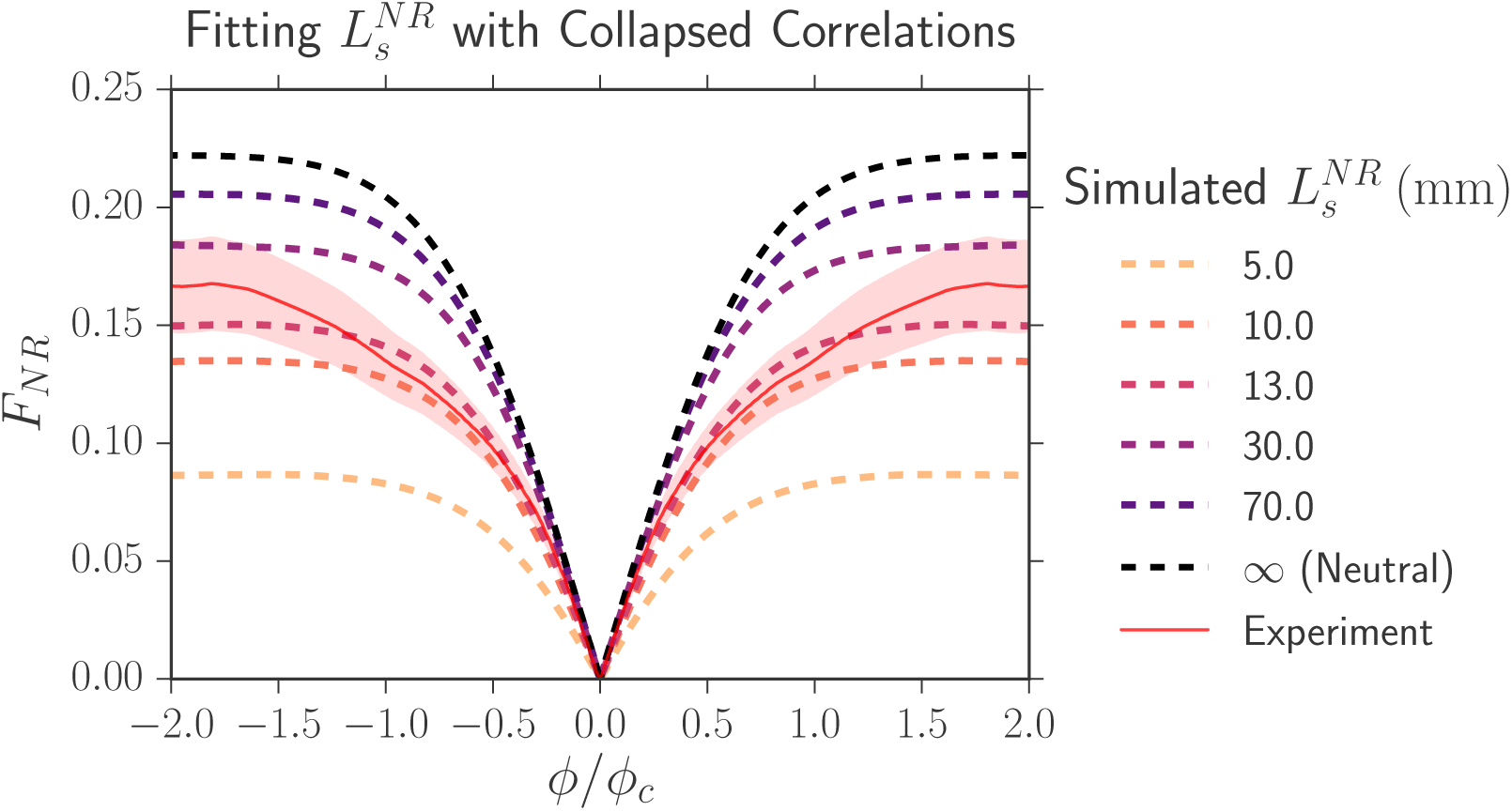
The collapsed two-point correlation function *F_NR_* between our eCFP/eYFP strain (*N*) and our mCherry strain (*R*), which were inoculated at fractions of 2/3 and 1/3, respectively, at a length expanded of *L* = 6.5 mm. The solid red line is experiment and the shaded region is the standard error of the mean. The dashed lines are the simulated universal correlation functions corresponding to different values of 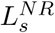. The best fitting selection length scale is 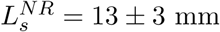

To determine the best-fitting value of *L_s_*, we calculated the sum of the squared displacements, weighted by the inverse of the experimental standard error squared, between experiment and simulation; the best-fitting *L_s_* was the one where the weighted sum of squares was minimized. To estimate the error in our fit, we assigned each potential value of *L_s_* a probability proportional to the inverse of the weighted sum of squares, normalized the probability distribution, and set the error in our fit of *L_s_* to the confidence intervals of the probability distribution.

To test that the resulting *L_s_* and *κ* could accurately predict the experimental dynamics at all *L* and not just the *L* where the correlation functions were fit, we plotted the experimental average fraction and correlation functions (solid lines) as we varied *L* and compared their values to those predicted by simulation (dashed lines) as shown in Figure 13. Figure 13 uses the same set of experimental data as that from Figure 12. The fit simulation always closely tracked the experimental values at all *L*, suggesting that our fitting technique was robust and could be used to describe the dynamics of our strains.

**Fig 13.**
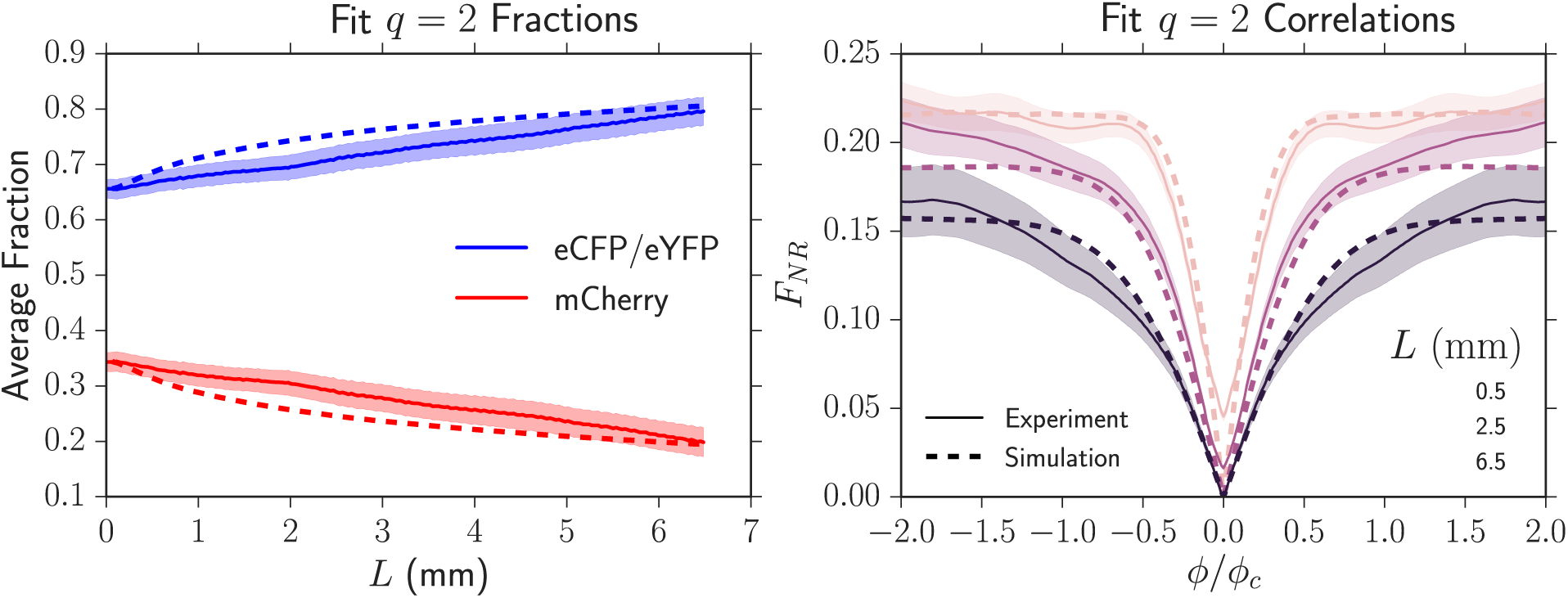
Experimental average fractions and two-point correlation functions (solid lines) and their predicted dynamics (dashed lines) by using the fit *L_s_* from Figure 12. The shaded region is the standard error of the mean. The simulated dynamics closely match the experimental dynamics, suggesting that our fitting technique to extract *L_s_* is robust and can be used to describe the dynamics of our strains at all *L*.

Competing pairwise combinations of strains and fitting their experimental two-point correlation functions with the simulated master curves allowed for accurate measurements of all 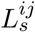 and *κ_ij_* as seen in Table 3. The *κ_ij_* are illustrated in Figure 7; smaller values of *κ_ij_* resulted in slower selective sweeps due to inflation. Although this technique was about a factor of 5 more precise than using the measured wall velocities 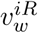 to determine 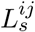, the upper bounds of the 95% confidence intervals were still very large as seen in Table 3; the potential values of *L_s_* had a very large tail.

**Table 3.** Results of fitting *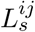* from the two-point correlation functions *F_ij_* and the resulting 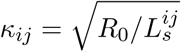 using the two-point correlation function technique. “CI” stands for confidence interval.

| Strain $i$ | Strain $j$ | $L_s^{ij}$ (mm) | 95% CI (mm) | $\kappa_{ij} = \sqrt{R_0/L_s^{ij}}$ |
| --- | --- | --- | --- | --- |
| eYFP/eCFP | mCherry | $13 \pm 3$ | [7, 38] | $0.52 \pm 0.07$ |
| Black | mCherry | $25 \pm 7$ | [14, 250] | $0.37 \pm 0.06$ |
| eYFP/eCFP | Black | $70 \pm 20$ | [30, 650] | $0.22 \pm 0.04$ |

#### Predicting the experimental dynamics

Having determined 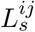 and *κ_ij_* from pairwise competitions between strains, we tested if we could predict the average fraction, the two-point correlation functions, and the annihilation asymmetry when the four *E. coli* strains were grown together (treating the eYFP and eCFP strains as neutral, so *q* = 3) in an independent experiment. We inoculated the four strains in equal proportions. Figure 14 shows experimental measurements of the average fractions and two-point correlation functions (solid lines) together with simulated predictions (dashed lines) that used the independently fit values; no additional fitting parameters were used. The predicted average fractions and correlation functions closely tracked the dynamics for *L* ≳ 3 mm. We attribute the deviations for *L* ≲ 3 mm to image analysis artifacts resulting from the presence of the black strain (see the Image Analysis section). At the largest length expanded of *L* = 6.5 mm where artifacts were minimal, the experiments matched the predictions within error. All average correlation functions were successfully predicted by the simulations; we only display *F_NR_* for simplicity.

**Fig 14.**
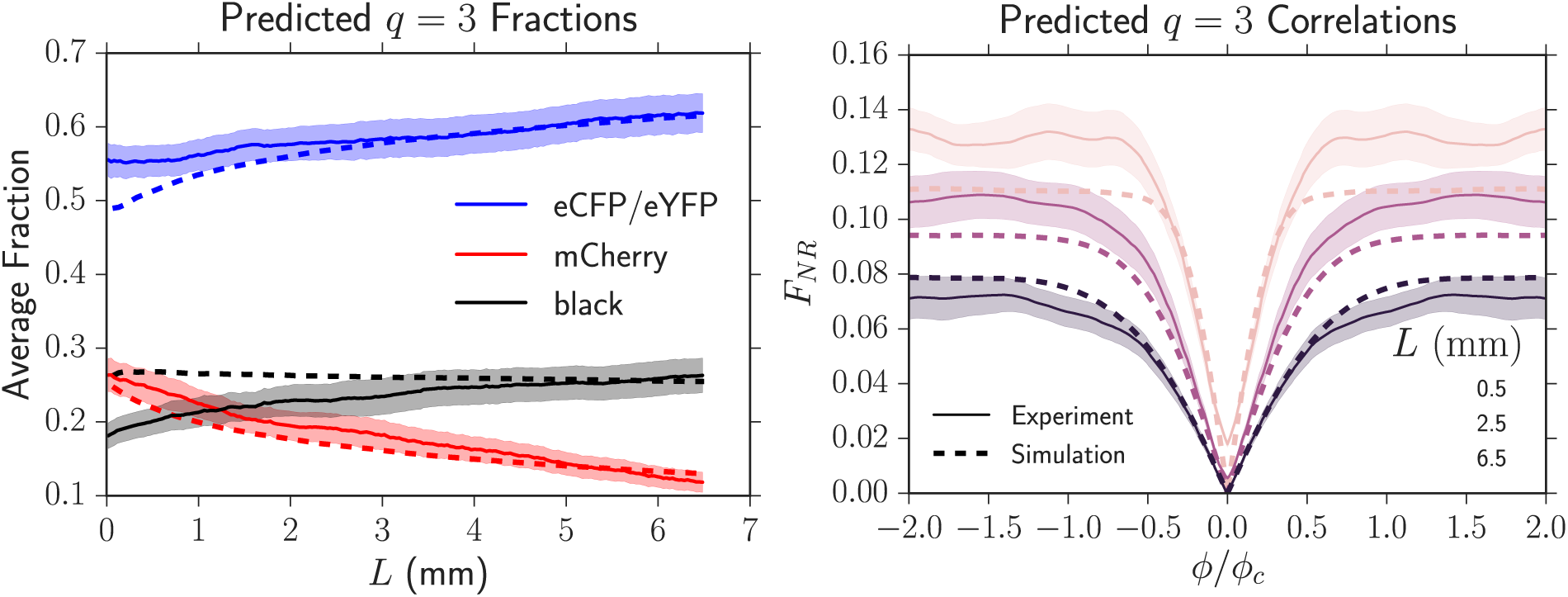
Experimental average fractions and two-point correlation functions (solid lines) of the four strains grown together with equal initial proportions and their predicted dynamics (dashed lines) from simulations using the set of *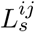*, *κ_ij_*, and *ϕ_c_* measured in independent experiments. No additional fitting parameters were used. The shaded region is the standard error of the mean. The simulated dynamics closely matched the experimental dynamics except at small lengths expanded (*L* ≲ 3 mm) where the black strain introduced significant image analysis artifacts.

In addition to predicting the average fractions and correlation functions, the fit values of 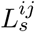 and *κ_ij_* predicted that the annihilation asymmetry would deviate only slightly from neutrality (at most a change of 0.1) over the length expanded by our colonies in every experiment, agreeing with our findings (Figure 10). This can be readily observed in Figure 11 which displays a simulation of two neutral strains (our eCFP and eYFP strains) and a less fit strain (our mCherry strain) inoculated in equal fractions. If we rescale the maximum distance expanded by our colonies of *L*_max_ = 6.5 mm by the smallest selection length (largest fitness difference) of 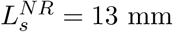, *L*_max_/*L_s_* ~ 0.5 and Δ*P* increases from 0 (neutrality) to at most 0.1. This small deviation from neutrality is within the uncertainty of our experimental measurement of Δ*P*. Evidently, certain quantities, like the average fraction and correlation functions, show signs of selection before others (in this case, the annihilation asymmetry).

The quantitative agreement between our model and our experiments suggests that the one-dimensional annihilating-coalescing random walk model can indeed be used to predict the dynamics of many competing strains with different fitnesses in a range expansion.

## Summary and conclusion

We investigated the evolutionary dynamics of four competing strains of *E. coli* in radial range expansions with differing selective advantages. We measured the average fraction *F_i_* of each strain, the two-point correlation functions *F_ij_* between strains, and the annihilation asymmetry Δ*P* with our image analysis toolkit [34]. Our simulations, which model the expansions as a one-dimensional line of annihilating and coalescing random walkers subject to deterministic drift on an inflating ring, showed us that these three quantities could be collapsed onto universal curves for a fixed set *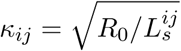* when the length expanded by the colony *L* was rescaled by any 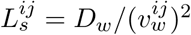 and the angular distance between strains *ϕ* was rescaled by 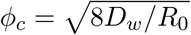.

To test if the random walk model could predict experimental dynamics, we independently calculated experimental values of 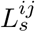, *κ_ij_*, and *ϕ_c_*, rescaled our experiments using the same procedure as our simulations, and compared the dynamics between the two. The simulations accurately predicted the dynamics of the average fraction, correlation functions, and annihilation asymmetry when all four of our strains were present with no additional fitting parameters. These results suggest that the random-walk model can act as a useful predictive tool when describing the evolutionary dynamics of range expansions composed of an arbitrary number of competing alleles with different fitnesses.

Along the way, we introduced a new technique that compared universal simulated correlation functions to experimental correlations to fit *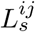*. The resulting values of *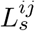* were about a factor of 5 more precise than directly evaluating 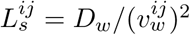 with the wall velocities extracted from the growth of sectors. Given our fit 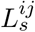, we evaluated 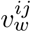 using 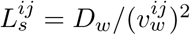 and the known *D_w_*; we compare these values to those extracted from single sectors in Table 4. The wall velocities from both measurements agreed within error, but the wall velocities obtained from our correlation method were at least a factor of two more precise than tracking single sectors. Although the correlation technique dramatically increased the precision in our evaluation of 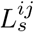, the resulting precision increase for 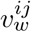 was less pronounced as 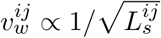. Nevertheless, it is clear that the correlation technique can be used to precisely extract small differences in fitness between spatially competing strains.

**Table 4.** Quantifying the wall velocities *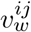* from our fits of *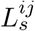* by using *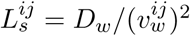* and an independently measured *D_w_* which was constant between experiments. The uncertainty in *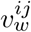* from the fit *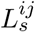* was calculated via error propagation.

| Strain $i$ | Strain $j$ | $L_s^{ij}$ (mm): fitting $F_{ij}$ | $v_w^{ij}$: from fit $L_s^{ij}$ | $v_w^{ij}$: tracking sectors |
| --- | --- | --- | --- | --- |
| eYFP/eCFP | mCherry | $13 \pm 3$ | $(9 \pm 1) \times 10^{-2}$ | $(6 \pm 2) \times 10^{-2}$ |
| Black | mCherry | $25 \pm 7$ | $(6 \pm 1) \times 10^{-2}$ | $(6 \pm 2) \times 10^{-2}$ |
| eYFP/eCFP | Black | $70 \pm 20$ | $(3.8 \pm 0.7) \times 10^{-2}$ | Undetectable |

Our work illustrates that the annihilating coalescing walk model can predict the experimental dynamics of an arbitrary number of competing alleles with different fitnesses in microbial range expansions. It is possible that this model could predict the dynamics of range expansions occurring outside of the laboratory. To predict the dynamics of expansions, however, the annihilating-coalescing walk model relies on a key set of parameters: the set of *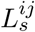*, the set of *κ_ij_*, and *ϕ_c_*. We found that the set of *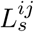* could *not* be predicted from the independent radial expansion velocities of our strains; standard techniques [17] using the relative ratio of expansion velocities to predict *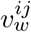*, and thus *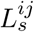*, yielded inconsistent results (see Appendix S2 where we quantify the discrepancy and postulate why it occurred). As the set of *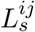* is so fundamental to the evolutionary dynamics of range expansions, future work should investigate why relative radial expansion velocities could not be used to accurately predict *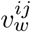* and thus *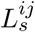* and whether this phenomenon is specific to *E. coli* range expansions or our specific strains. It would also be interesting to incorporate the reported super-diffusive motion of domain walls [8, 9] into our simplified simulations and theoretical analysis. The random walk model’s ability to successfully predict the evolutionary dynamics of our experiments suggests that annihilating and coalescing genetic domain walls subject to diffusion and selection-induced displacement provide a useful conceptual framework from which to understand range expansion dynamics.

## Supporting Information

**S1 Appendix. Supplemental theory.**

**S2 Appendix. Quantifying the discrepancy between radial expansion velocity and wall velocity.**

**S1 Fig. Image processing artifacts introduced by using a non-fluorescent (i.e. black) strain.** To estimate the image analysis artifacts introduced by using a non-fluorescent, black strain we performed an experiment with three fluorescent strains (eCFP, eYFP, and mCherry in equal initial proportions) and analyzed the data twice: once where we included all three fluorescent channels and once where we excluded the eCFP channel and treated it as if it were a black strain. We compared the black-substituted average fractions *F_i_* (the dashed lines) to the real fractions as a function of radius (the solid lines). At a small radius relative to *R*_0_ = 3.5 mm, the error from introducing a black strain was large; this is likely because we defined black as the absence of any other channels and channels typically had large overlaps close to the homeland. At large radius, the error from introducing a black strain was negligible.

**S2 Fig. Determining *R*_0_.** To fit the radius *R*_0_ where our image analysis package became accurate, we inoculated 80% of mCherry, 10% of eCFP, and 10% of eYFP in 10 range expansions and tabulated the average fraction of each strain. The inoculated fractions are illustrated by dashed lines. As seen in the plot, at a radius of approximately *R*_0_ = 3.50 ± 0.05 mm the average fractions approximately matched the initial inoculated fractions. Our image analysis package inaccurately predicted fractions in the homeland because of significant overlap between the strains.

**S3 Fig. Error bars when fitting *D_w_*.** The same as the right side of Figure 5 except with error bars; the shaded areas are the standard error of the mean.

**S4 Fig. Average cumulative annihilations and coalescences for two, three, and four strains.** All strains were inoculated in equal fractions except for the experiment with 10% of eCFP, 10% of eYFP and 80% of mCherry. The annihilation and coalescence rates (the slope of the respective curves) decrease as radius increases as there are less domain walls due to previous collisions and also because inflation decreases the probability of two walls colliding per length expanded. As the number of colors increases, coalescences occur more often than annihilations.

**S5 Fig Confirming simulation accuracy.** We simulated a single fit sector sweeping through a less fit strain. It is expected that the fit strain sector dynamics satisfy 〈*ϕ* − *ϕ*_0_〉 = 2*v_w_* ln (*R*/*R*_0_) and Var (*ϕ*) = 4*D_w_* (1/*R*_0_ − 1/*R*), as seen in Appendix S1. To test that our simulation appropriately reproduced this behavior, we quantified the average angular growth 〈*ϕ* − *ϕ*_0_〉 and angular variance Var(*ϕ*) as we varied the simulation parameters *N*_0_ (initial number of cells), *r* (selective advantage of the fitter strain), and *d* (distance the colony expanded each generation). The cell width, *a*, was kept constant. These parameters relate to the sector dynamics via *D_w_* = *a*^2^/(2*d*), *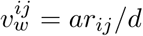*, and *R*_0_ = (*N*_0_*a*)/(2*π*). We confirmed that both the average angular growth 〈*ϕ* − *ϕ*_0_〉 and angular variance Var(*ϕ*) had the correct functional form and dependence on the microscopic parameters (the dashed black line). In the main text, we used *d* = *a* for simplicity.

**S6 Fig. Collapsed average fraction and annihilation asymmetry on a linear scale.** Identical to Figure 11 except the *y*-axis of *F*(*L*/*L_s_*, *κ*) is placed on a linear scale, which may be useful for comparison with experiments.

**S7 Fig. Collapse of *F_ij_*.** We ran four simulations where we varied *v_w_* (the velocity that the two other more fit strains swept through the less fit strain) and *R*_0_ such that *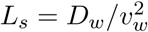*changed but *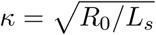* was fixed. Each simulation has a different symbol in the plot. We found that *F_ij_* could be collapsed at the same *L*/*L_s_* as long as *κ* remained fixed (we arbitrarily set it to *κ* = 0.4) and as long as the angular variable *ϕ* was rescaled by 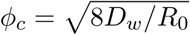. If *ϕ_c_* approached the system size *ϕ_c_* ≈ 2*π*, *F_ij_* could not be collapsed onto the above curves due to finite size effects. Note that even though we only show *F_RR_*, all correlation functions *F_ij_* could be collapsed using this procedure.

**S8 Fig. Collapse of average fraction and annihilation asymmetry.** We simulated three competing strains and arbitrarily chose a fixed set of *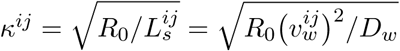* to reflect differing selective advantages between the strains. We set the initial radius of the expansion *R*_0_ = *N*_0_*a*/(2*π*) to *N*_0_ = {200, 2000, 20000} in three simulations and altered 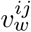 to tune the set of *κ^ij^*’s to their fixed values. We found that the dynamics collapsed as long as *L* was rescaled by any selection length scale in the system, i.e. 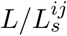 (we chose to use 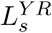). The diamond simulation, corresponding to *N*_0_ = 200, deviated slightly from the other simulations because of finite size effects (i.e. when *ϕ_c_* ~ 2*π*).

## Acknowledgments

BTW would like to thank Matti Gralka, Paula Villa Martin, Miguel A Muñoz, Severine Atis, Markus F. Weber, Kirill Korolev, and Steven Weinstein for helpful discussion and advice. Research by BTW is supported by the Department of Energy Office of Science Graduate Fellowship Program (DOE SCGF), made possible in part by the American Recovery and Reinvestment Act of 2009, administered by ORISE-ORAU under contract no. DE-AC05-06OR23100, by the US Department of Energy (DOE) under Grant No. DE-FG02-87ER40328, as well as Harvard University’s Institute for Applied Computational Science (IACS) Student Fellowship. BTW and DRN benefitted from support from the Human Frontiers Science Program Grant RGP0041/2014 and from the National Science Foundation, through grants DMR1608501 and via the Harvard Materials Science and Engineering Center, through grant DMR1435999. MOL acknowledges support from NSF grant DMR-1262047, the UPenn MRSEC under Award No. NSF-DMR-1120901, the US Department of Energy, Office of Basic Energy Sciences, Division of Materials Sciences and Engineering under Grant No. DE-FG02-05ER46199, and from the Simons Foundation for the collaboration “Cracking the Glass Problem” (Grant No. 454945).

## Appendix S1 Supplemental theory

As discussed in the introduction, it is difficult to make analytical progress when modeling a many-allele range expansion as a line of annihilating and coalescing domain walls subject to diffusion and deterministic, selection-induced motion. This is because the moment hierarchy of an equivalent system, the *q*-color Voter Model, does not close [1]. Much is understood, however, about the neutral dynamics of many annihilating and coalescing walls; analytical predictions exist for spatial correlation functions [2] and relative annihilation and coalescence rates [2–4]. In addition, results for the dynamics of a monoclonal single sector (bordered by domain walls) of a more fit strain sweeping through a less fit strain are available [1, 5, 6]. In this section, we review previous theoretical work and introduce theoretical advances relevant to the main text.

### The Stepping-Stone and Voter models with selection

The population dynamics of range expansions with flat fronts can be modeled as the evolution of a one-dimensional line of individuals subject to growth/replication, death, and diffusion with the constraint of a constant population density [1]. Two well-studied microscopic models of the dynamics are the Voter model [6] and the Stepping Stone model [7]. The Voter model is equivalent to a one-dimensional *q*-state Potts model [8, 9] governed by zero-temperature Glauber dynamics [10]; individuals in the population are replaced by one of their neighbors with a constant probability per time. The stepping stone model (prior to taking a continuum limit) assumes that there are many connected “islands” that individuals diffuse to and from governed by Moran reproduction [11]; each of these islands has a population size of *N*. The Voter model can be viewed as a stepping stone model where the local population of each island is *N* = 1.

When either model is coarse-grained in space, the resulting stochastic differential equation governing the evolution of 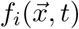, the fraction of strain *i* at position 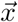 at time *t*, is the same, but with different values for the model parameters and boundary conditions. The stochastic differential equation for an arbitrary number of competing strains along the line is

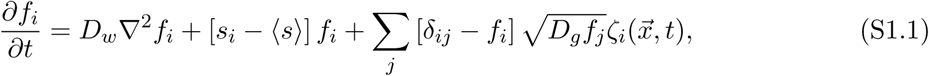

which is a spatial generalization of the equation studied by Good et al. [12]. Here, *D_w_* is the spatial diffusion coefficient of each strain and is the same *D_w_* used in the main text, *s_i_* is the dimensionless fitness of strain *i*, 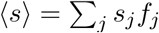 is the mean fitness of all strains locally, and *D_g_* parameterizes genetic drift. The function 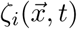 is a white noise random variable with the properties 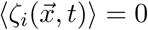 and 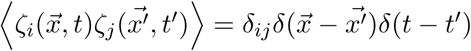, where these noises are interpreted in the Itô sense [13]. The noise term on the far–right captures fluctuations due to the limited population size. Equation (S1.1) reduces to standard equations in the literature [1] when describing with two competing strains. Unfortunately, equation (S1.1) becomes analytically intractable when multiple strains and selective advantages are present because of the closure problem:*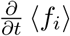* depends on 〈*f_i_ f_j_*〉, and *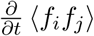* depends on 〈*f_i_ f_j_ f_k_*〉, etc. The moment hierarchy does not close.

### Neutral correlation functions

Much is known about equation (S1.1) in the neutral case where all *s_i_* = 0 as discussed in the main text [1]. Unsurprisingly, it can be shown that the average fraction *F_i_* = 〈 *f_i_* 〉 equals the initial inoculated fraction *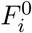* at all times assuming the initial inoculant is uncorrelated where the bracket indicates an average over many expansions. Although the average fraction is constant in the neutral case, the two–point correlation function broadens due to the coarsening of genetic domains. Upon adopting polar coordinates for radial expansions and letting *L* = *ut* = *R* − *R*_0_ where *u* is the radial expansion velocity, it can be shown, using equation (S1.1) when all *s* = 0, that the dynamics of the average two-point correlation functions *F_ij_*(*ϕ*, *L*) = 〈*f_i_ f_j_*〉 are governed by (in polar coordinates),

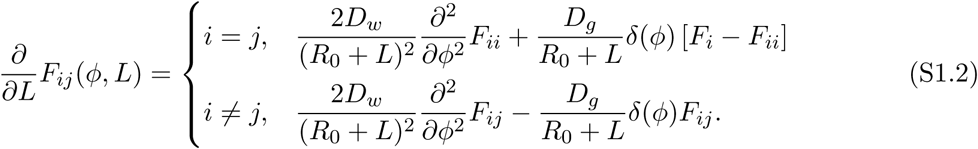

where *ϕ* is the angular distance between points at the frontier. For the Voter model with deme size *N* = 1, the boundary conditions are given by 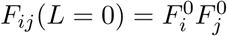,*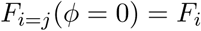*,and 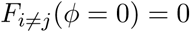; these conditions make the delta functions *δ*(*ϕ*) disappear. *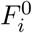* and 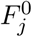 are the initial inoculated fractions of strains *i* and *j*. Solving these equations by making a “conformal time” substitution [6, 14, 15] yields

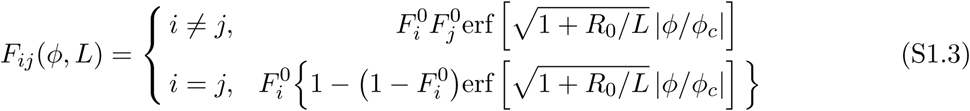

where the characteristic angular width of *F_ij_* is given by 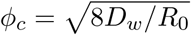; again note that *D_w_* is the same as that from the main text. Figure S1.1 contains plots of *F_ij_* (*ϕ*, *L*) for both *i* ≠ *j* and *i* = *j*. As *L* → ∞, *F_ij_* approaches a constant shape given by the error function because inflation will eventually completely dominate the diffusive motion of boundary walls which brings coarsening of genetic domains to a halt. When *ϕ* ≫ *ϕ_c_*, we have 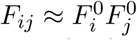, because the different genetic regions become uncorrelated. Note that if *ϕ_c_* approaches 2*π*, this limit is impossible to achieve and the correlation function will not factorize. These neutral results for *F_ij_*(*ϕ*, *L*) tabulated above can be used as a null model when we introduce selection.

**Fig S1.1.**
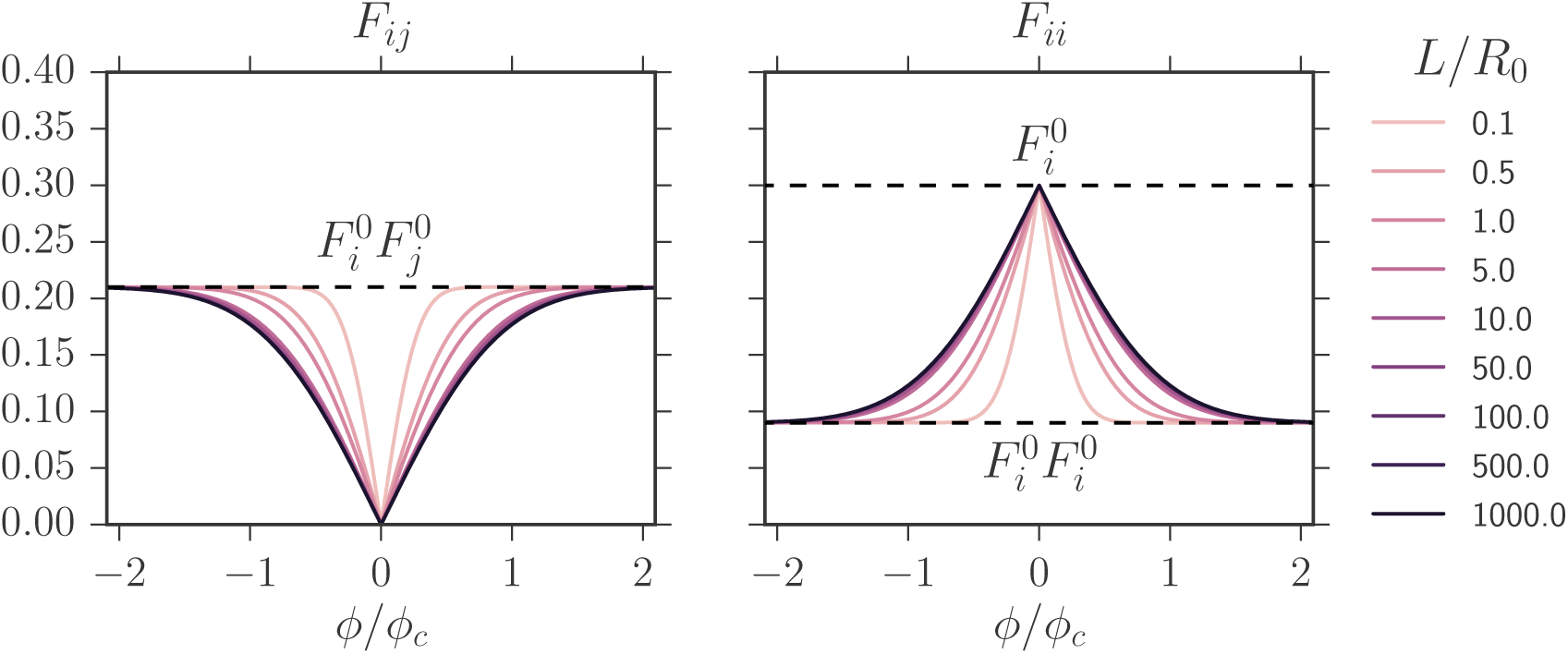
Voter model predictions for *F_ij_*(*ϕ*, *L*) from eq. (S1.3) for *i* ≠ *j* on the left and *i* = *j* on the right. *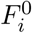* and *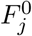*, the initial inoculated fractions of strains *i* and *j*, were set to 0.3 and 0.7 respectively. The product 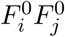 determines the asymptote of the correlation function for large angular separation. Note that *L*/*R*_0_ ≥ 0.

From the *F_ij_*(*ϕ*, *L*) above, we define the heterozygosity correlation function as [1]

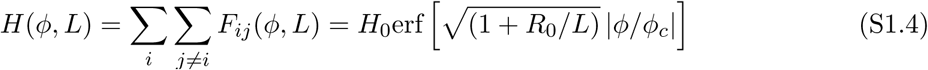

where *H*_0_ is the heterozygosity when *L* = 0 (this assumes that that the initial condition is uncorrelated), or 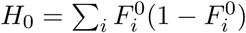. The heterozygosity can be thought of as the probability that two points separated by an angle of *ϕ* at a length expanded of *L* are occupied by different strains; it is a spatial measure of genetic diversity. This result is used to determine the wall diffusion constant *D_w_* in the Measuring *D_w_* section in the main text.

### Neutral annihilation and coalescence probabilities

Upon collision, the diffusing domain walls either annihilate or coalesce as illustrated in Figure 1 of the main text. Upon regarding these genetic boundaries as world lines of chemical species, these processes can be described using the language of chemical reactions,
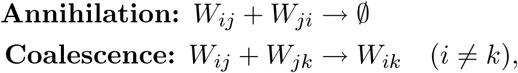

where *W_ij_* is a domain wall such that the strain on the left is of type *i* and the strain on the right is of type *j*. Note that the inner indices of colliding domain walls are always identical, because two neighboring domain walls always have a common strain between them.

If an arbitrary integer number of *q* neutral colors are inoculated at random locations with *equal* initial proportions on a one-dimensional lattice, it is known that the probability of obtaining an annihilation per domain wall collision is given by *P_A_* = 1/(*q* − 1) and the probability of obtaining a coalescence per collision is given by *P_C_* = (*q* − 2)/(*q* − 1) [2–4].

To determine how unequal global fractions of each neutral strain alters *P_A_* and *P_C_*, we write *P_A_* and *P_C_* in terms of *P_ijk_*, the probability that a collision between domain walls *W_ij_* and *W_jk_* occurred per collision:

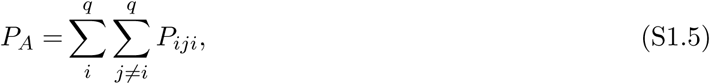

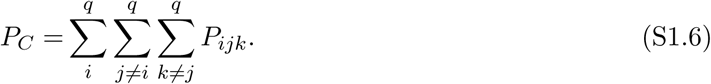

We expect that for *q* neutral colors, the chance of a particular color combination in a collision *ijk* with color *i* on the left and color *k* on the right and *j* in the middle to be proportional to the product 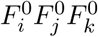 of the initial color fractions.We therefore expect that

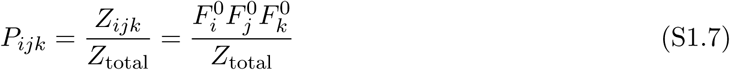

where *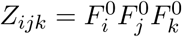* and the normalization constant is *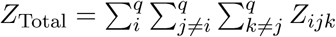*, i.e.the sum of all *Z_ijk_*.

Using the simulations described in the Simulation section, we checked eq. (S1.7) for *q* = 3 neutral strains. The left side of Figure S1.2 displays the simulated values of *P_ijk_* and our theoretical predictions for three neutral strains inoculated with initial fractions {*F*_1_ = 0.1, *F*_2_ = 0.3, *F*_3_ = 0.6} in a linear range expansion; our theoretical predictions, represented by black dashed lines, match the simulation results. As our predictions for *P_ijk_* were correct, our predictions for *P_A_* and *P_C_* were also correct as they were composed of sums of *P_ijk_*. Inflating simulations with the same *F_i_* also returned the same values of *P_ijk_* and thus *P_A_* and *P_C_*. Inflation changes the rate at which annihilations and coalescences occur, but not their relative proportions.

**Fig S1.2.**
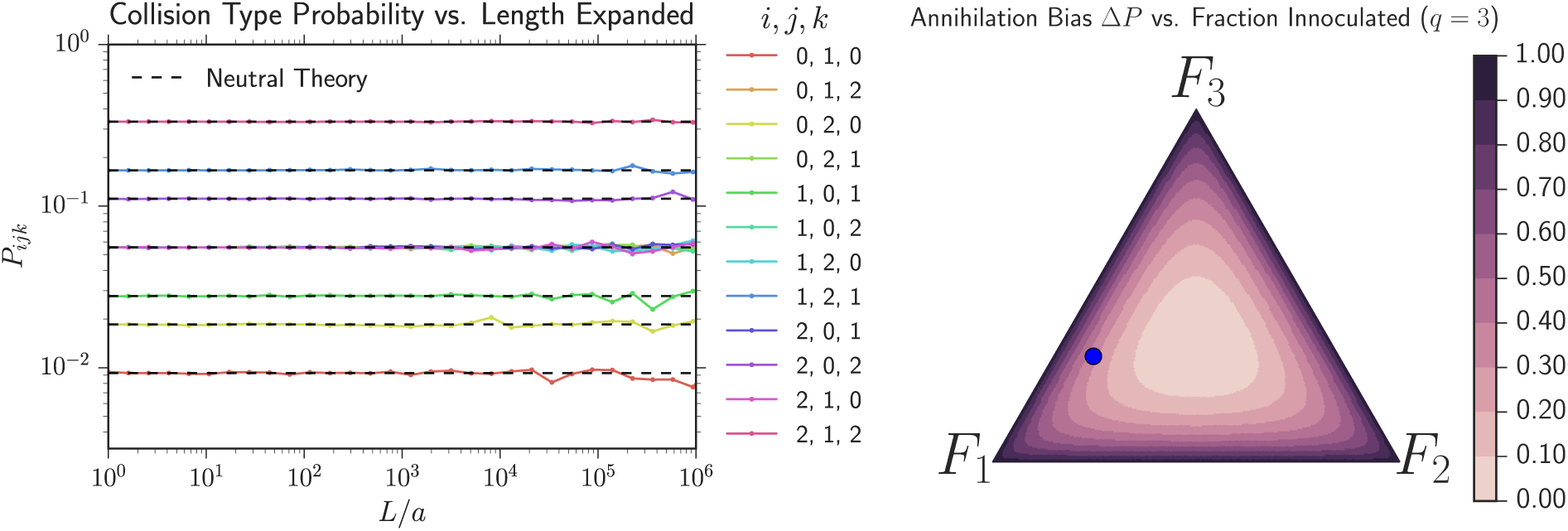
*Left*: Probability *P_ijk_* of a domain wall collision with color *i* to the left of the walls, color *j* between the walls, and color *k* to the right vs. length expanded. We simulated *q* = 3 neutral strains with initial fractions {*F*_1_ = 0.1, *F*_2_ = 0.3, *F*_3_ = 0.6} in a linear expansion, averaged the results of 1000 simulations, and calculated *P_ijk_* (solid lines) and compared its value to that from eq. (S1.7) (dashed black lines). *L*/*a* is the length expanded divided by the cell size and is equivalent to the elapsed time in generations. The simulation confirms the predictions of eq. (S1.7). *Right:* The annihilation bias Δ*P* = *P_A_* − *P_C_*, where *P_A_* and *P_C_* are the probabilities of obtaining an annihilation or coalescence per domain wall collision respectively, calculated via eqs. (S1.5) and (S1.6) as a function of initial inoculated fractions for *q* = 3 neutral colors. Δ*P* is independent of length expanded for neutral strains. The large blue dot corresponds to the initial conditions that were used on the left. Δ*P* assumes its minimum value Δ*P* = 0 when *q* = 3 colors are inoculated in equal fractions and is maximized on the boundaries of the ternary diagram corresponding to Δ*P* = 1. Discrete colors were used to more clearly highlight the contours of Δ*P*.

As discussed in the main text, to efficiently quantify the difference between the annihilation and coalescence probabilities, we defined the “annihilation asymmetry” Δ*P*(*L*) = *P_A_*(*L*) − *P_C_*(*L*) as the difference in probability of obtaining an annihilation vs. a coalescence in a given collision at a distance expanded of *L*. In the neutral case, Δ*P* is independent of *L*. The right side of Figure S1.2 displays a ternary diagram illustrating all possible values of Δ*P* that can be reached when inoculating *q* = 3 neutral colors in different proportions. The blue dot corresponds to the initial conditions where the *P_ijk_* probabilities were calculated in the plot to its left. For a combination of *q* colors present in an expansion, Δ*P* is minimized when *q* colors are inoculated in equal fractions and is maximized when one of the fractions of the *q* colors goes to zero.

#### Single sector dynamics with selection

We first review a simple phenomenological model [5, 6, 16] of the width *w* of a single sector of a more fit strain sweeping through a less fit strain incorporating both wall diffusion and deterministic motion due to selective differences. Let *x* be the position of one of the domain walls of a sector. We quantify a domain wall’s displacement *dx* over a a front expansion distance of *dL* by the parameters 2*D_w_* = *d*Var(*x*)/*dL* (Var(*x*) = 〈*x*^2^〉 − 〈*x*〉^2^ is the variance), describing the diffusive motion of the wall, and *v_w_* = *d*〈*x*〉/*dL*, describing its deterministic motion, as discussed in the Introduction and illustrated in Figure 1 of the main text.

We first describe a linear range expansion and then extend our treatment to a radially inflating expansion. If we track the distance *w* between two walls that are sweeping through a a less fit strain per length expanded *L*, as sketched on the right side of Figure 1 of the main text, the dynamics of *w* is given by

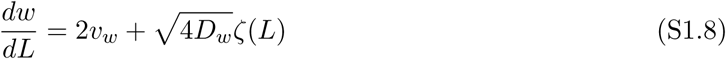

where *ζ*(*L*) is white noise with 〈*ζ*(*L*)〉 = 0 and 〈*ζ*(*L*)*ζ*(*L*′)〉 = *δ*(*L* − *L*′) and should be interpreted in the Itô sense [13]. The factors of 2 in front of *v_w_* and 4 in front of *D_w_* arise because we monitor the distance between *two* domain walls. Note that we make the smooth front approximation that neglects the roughness of the expansion boundary, forcing *v_w_* to be constant perpendicular to the growth direction [17] and forcing domain wall motion to be diffusive [6]. Within this approach, diffusive effects scale as 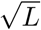 while deterministic effects scale as *L*; hence, at short expansion distances, diffusion dominates the sector width while at larger length scales, deterministic motion becomes apparent. A crossover length scale *L_s_* follows by equating the deterministic average displacement (from the first term of (S1.8)) with the root mean squared displacement associated with the second term,

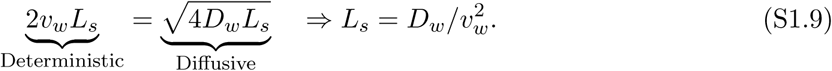

*L_s_* is the distance that a linear expansion must expand in order for selection to become dominant over diffusive effects [1, 6, 18].

What changes in the radially expanding case? We now shift to radial coordinates. Upon setting *L* = *R* − *R*_0_ where *R*_0_ is the radius at which the alleles first fix, and denoting the angular width between the two sectors as *ϕ* = *w*/*R*, our phenomenological stochastic model becomes [6]

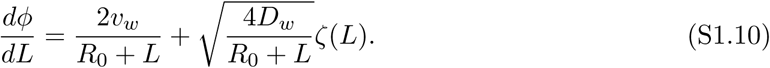

The mean and variance of the sector width *ϕ* are, with *R* = *L* + *R*_0_,

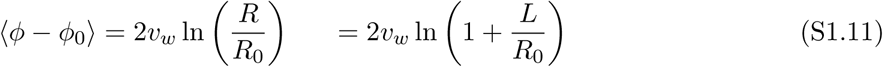

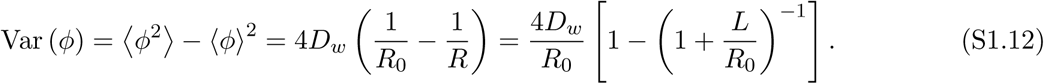

Eq. (S1.11) describes how the boundaries of the more fit domain sweep out a logarithmic spiral as the expansion inflates [6, 16, 17], and eq. (S1.12) shows that the effective angular diffusion constant decreases as the radius *R* = *R*_0_ + *L* increases. If one now equates the deterministic displacement of the boundaries to diffusive effects, in analogy with equation (S1.9), we find that the crossover between diffusive wandering of the sector width and a deterministic logarithmic sweep occurs at an expansion distance *L_I_* that satisfies

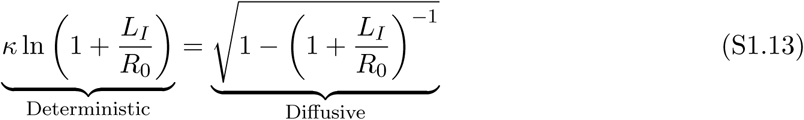

where the dimensionless parameter *κ* is an inflationary selective advantage, [6] *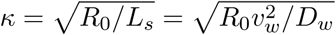* and *L_I_* is the inflationary analog of *L_s_*, the length scale at which selection dominates over diffusion on an inflating boundary [6, 16]. Fig. 7 of the main text displays the numerical solution of eq. (S1.13) for *L_I_*(*κ*).

The impact of *κ* on domain behavior is demonstrated visually in Figure S1.3. Three simulations were conducted utilizing the algorithm from the Simulation Section where a more fit yellow strain swept through a less fit red strain. *L_s_* was kept constant but *κ* was varied by altering the initial radius of each expansion. As *κ* decreased, inflation played a larger role and dramatically slowed down the sweep of the more fit strain.

**Fig S1.3.**
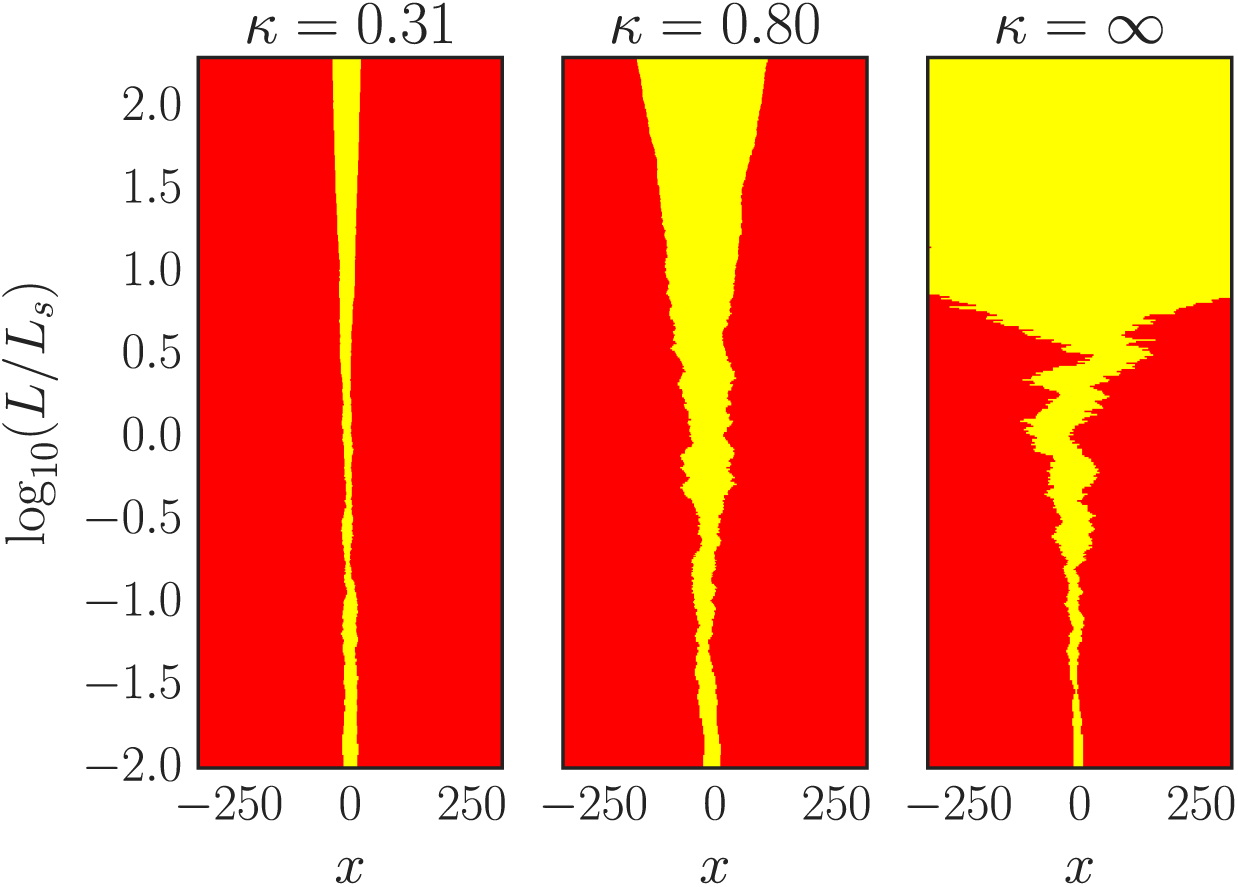
The impact of *κ* on domain behavior. Three simulations were conducted utilizing the algorithm from the Simulation section where a more fit yellow strain, initially occupying a width of 10 cells at the front (the horizontal axis is in units of cell widths), swept through a less fit red strain. In our simulations, we kept the radius of expansions fixed at *R*_0_ and accounted for inflation by appropriately decreasing the “jump size” of domain walls (see the Simulation Section); this leads to the identification that *x* = *R*_0_*ϕ* where *ϕ* is the angular position along the radially expanding front. As *κ* decreases from right to left, inflation plays a larger role and dramatically slows down the distance swept by the more fit strain due to the decreasing domain wall jump length. *κ* = ∞ was obtained by not inflating the domain (a linear range expansion) with periodic boundary conditions; the expansion proceeds along a cylinder.

## Appendix S2 Quantifying the discrepancy between expansion velocity and wall velocity

In our experiments, we observed that the the eCFP and and eYFP strains expanded faster than the black strain which in turn expanded faster than the mCherry strain (see Table 1). In addition, we found that the eCFP and eYFP strains swept through the black strain which swept through mCherry when competing in the same expansion (see Table 4). These observations are consistent with a picture in which a larger expansion velocity difference leads to a larger wall velocity. Korolev et al. studied the connection between radial expansion velocity and wall velocity in detail for *S. cerevisiae*. Using geometric arguments, they argued that if the front of an expansion is sufficiently smooth, a domain wall bordering a strain *i* and a less fit strain *j* will have a constant wall velocity 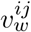 towards the less fit strain dependent only on the ratio of radial expansion velocities *u_i_*/*u_j_* [1, 2] given by

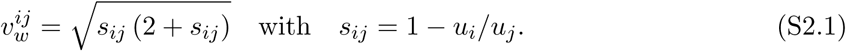

Korolev et al. found that this relationship holds in *S. cerevisiae* expansions at large lengths expanded; at small lengths expanded, the prediction overestimates the wall velocity [1].

We tested if the average radial expansion velocities *u_i_* of our *E. coli* strains could predict 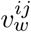 via eq. (S2.1) (which in turn could be used to evaluate 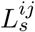), as it is easier for experimentalists to measure radial expansion velocities than 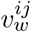 directly. We measured *u_i_* using the procedure described in the Characterizing radial expansion velocities section and quantified *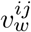* by directly tracking the growth of single sectors of a faster expanding strain sweeping through a slower expanding (less fit) strain. Characteristic single sectors of each strain sweeping through mCherry can be seen in Figure 6 in the main text. For a given wall velocity in radial expansions, the domain walls of more fit strains will, on average, sweep logarithmic spirals through less fit strains. Logarithmic spirals have been observed in yeast expansions at large lengths expanded [1]. Specifically, the average angular width of a sector of strain *i* sweeping through strain *j* is given by (see Appendix S1 for more details)

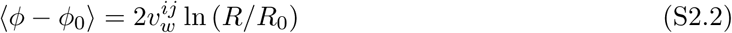

where *ϕ*_0_ is the initial angular width of the domain at *R*_0_ and *ϕ* is the width at radius *R*. 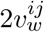 can thus be extracted from the slope of a linear regression fit of 〈*ϕ* − *ϕ*_0_〉 vs. ln(*R*/*R*_0_).

Although the rank order of our strains’ expansion velocities was constant between plates, the precise value of the velocities varied. To control for this fact, we grew the colonies used to determine *u_i_* and *u_j_* on the same plate as the colony used for evaluating 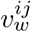 directly from the motion of domain walls, i.e., on one plate we inoculated a colony of a fast growing strain *i*, a colony of a slower growing strain *j*, and a mixed colony composed of 10% of strain *i* and 90% of strain *j*. The mixed colony’s ratio of strains was chosen so that single sectors of the more fit strain would form.

Using the radial expansion velocities of the *i* and *j* expansions (*u_i_* and *u_j_*) in eq. (S2.1), we computed a predicted 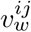 which was then used to predict the average sector width 〈*ϕ* − *ϕ*_0_〉 via eq. (S2.2). The comparison between the predicted average sector width and the experimental average sector width is shown in Figure S2.1. We found that for all 10 replicates for each pairwise competition of strains, *s_ij_* = 1 − *u_i_*/*u_j_* overestimated the magnitude of 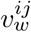 by almost a factor of 5; the predicted mean angular width from eq. (S2.2) overestimated our experimental measurements by over 3 standard deviations at the largest length expanded.

**Fig S2.1.**
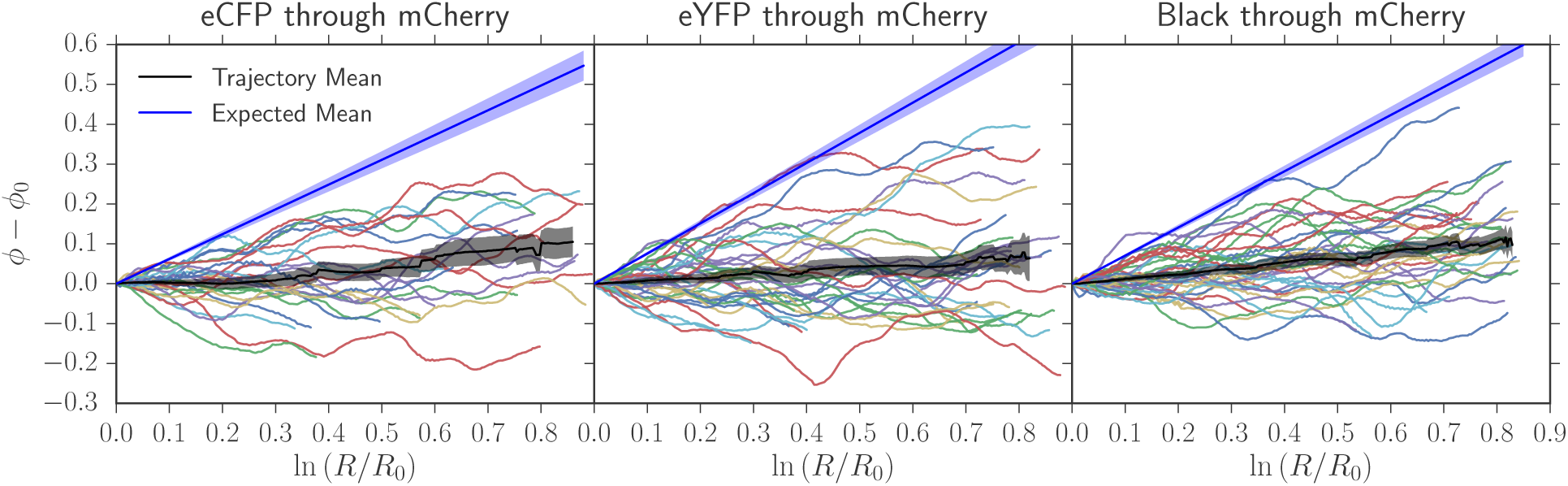
Expected average angular growth of sectors (blue) from equation (S2.2) using strains’ relative expansion velocities vs. the actual average angular growth (black). The shaded areas are the standard error of the mean and the colored lines are individual traces of sectors’ angular width. Equation (S2.2), using the predicted wall velocity 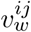 extracted from the ratio of the strain expansion velocities in eq. (S2.1), overestimated the average angular width at the largest ln(*R*/*R*_0_) by over 3 standard deviations.

We also compared the predicted values of *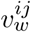* to those calculated from the experimental slope of the average angular growth of sectors (see eq. (S2.2)) in Table S2.1. We found the value of every experimental wall velocity to be 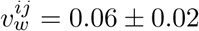; the ratio of velocities predicted the wall velocities to be on the order of *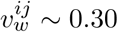*.

**Table S2.1.** Predicted wall velocity *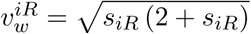* from the ratio of expansion velocities *s_iR_* = *u_i_*/*u_R_* − 1 vs. direct measurements of *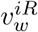*by tracking the angular growth of many sectors of a more fit strain sweeping through mCherry. Our predictions of the wall velocity overestimated the actual wall velocity by abogut a factor of 5. Figure S2.1 displays the single sector trajectories that were averaged to extract *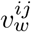*.

| Sweeper | $v_w^{iR} = \sqrt{s_{iR}(2 + s_{iR})}$ | $v_w^{iR}$: tracking sectors |
| --- | --- | --- |
| eYFP | $0.34 \pm 0.03$ | $0.06 \pm 0.02$ |
| eCFP | $0.32 \pm 0.04$ | $0.06 \pm 0.02$ |
| Black | $0.26 \pm 0.04$ | $0.06 \pm 0.02$ |

Both Figure S2.1, displaying the predicted average angular width and our experimentally measured average width, and Table S2.1, where we compared the predicted value of 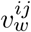 to its experimentally measured values, indicate that individual expansions velocities overestimate the wall velocities observed when using equations (S2.1) and (S2.2). Note that when a mixed colony was grown alone on a plate and *s_ij_* was predicted from many replicates of pure strains of *i* and *j* grown on separate plates made on the same day, the differences in radial expansion velocities still overestimated *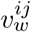* by a factor of 5 to 10; the presence of three colonies on a plate did not change our conclusion.

As mentioned above, Korolev et al. [1] found that geometric predictions overestimated the wall velocity in yeast expansions for small lengths expanded. However, at large lengths, the wall velocity approached its predicted value. A similar effect is likely occurring with our *E. coli* strains, except that we do not find an approach to the predicted wall velocity value over the length of our experiments; the wall velocity was *always* less than the prediction of equation (S2.1). It is possible that unaccounted mechanical forces, such as surface or line tensions, damp the ability of more fit strains to bulge outwards, preventing geometric arguments from applying. Another explanation is that simple geometric arguments describing wall motion no longer hold as a colony roughens. There is also the possibility of unexpected mutualistic or antagonistic chemical secretions between strains. However, this explanation seems unlikely because the ratio of expansion velocities *s_ij_* of the three colonies grown on the same plate matched the *s_ij_* of strains grown on independent plates (when using the same batch of plates). Order of magnitude estimates of diffusion constants suggest that mutualistic or antagonistic secretions would have diffused over the entire plate during the 8 days of an experiment and would have likely changed the relative fitnesses of the expanding colonies.

In conclusion, for the *E. coli* strains and the growth conditions we used, it was not possible to predict the wall velocities from independently measured radial expansion velocities using the geometrical argument underlying eq. (S2.1). Understanding the origin of this discrepancy is an interesting avenue for future investigation. Since this work focuses on competition within colonies, we use directly measured wall velocities from both image analysis (see the Measuring *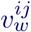* section) and our two-point correlation function fitting technique (see Table 4) to predict our experiments’ evolutionary dynamics.

## Supplementary Figures

**Fig S1.**
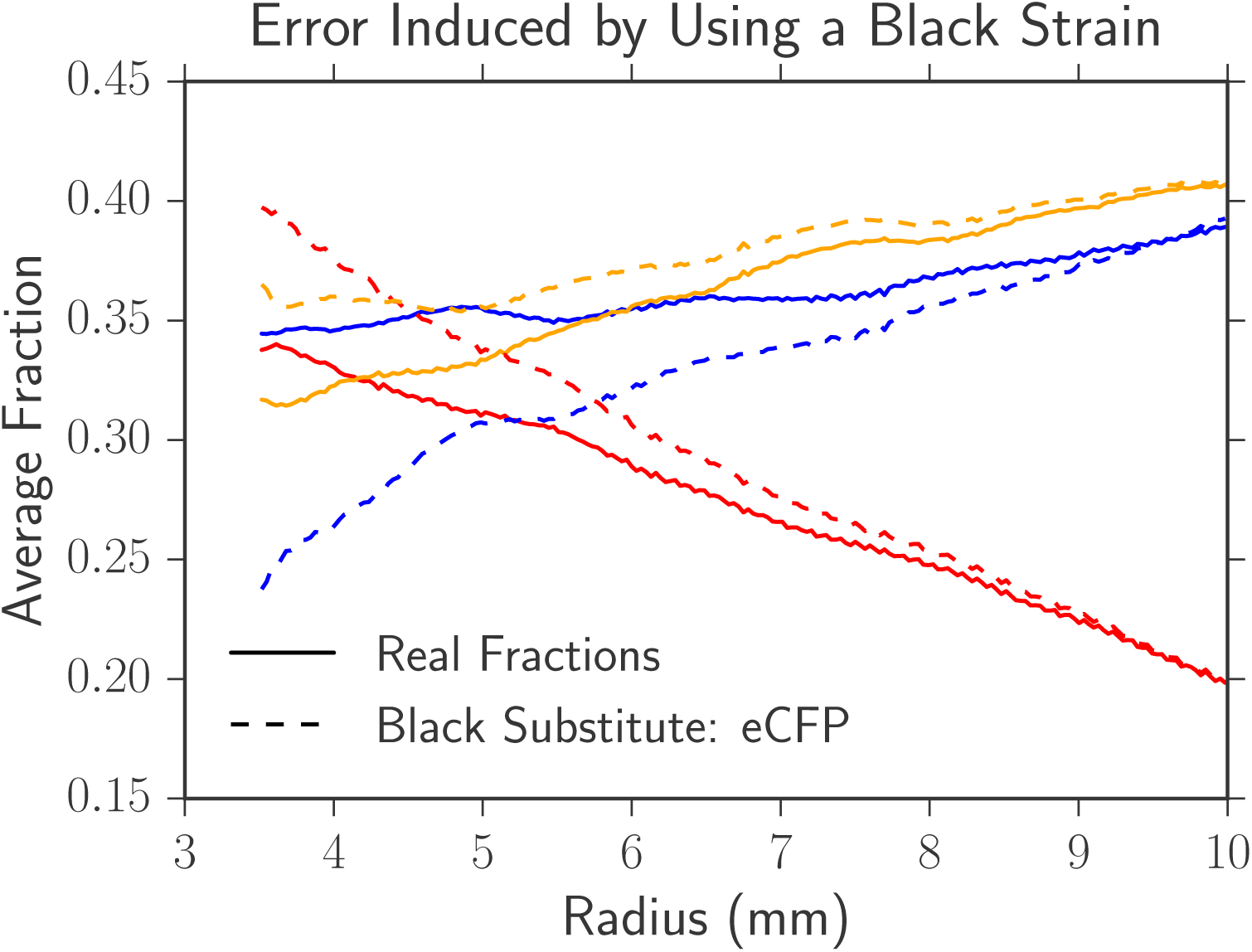
Image processing artifacts introduced by using a non-fluorescent (i.e. black) strain. To estimate the image analysis artifacts introduced by using a non-fluorescent, black strain we performed an experiment with three fluorescent strains (eCFP, eYFP, and mCherry in equal initial proportions) and analyzed the data twice: once where we included all three fluorescent channels and once where we excluded the eCFP channel and treated it as if it were a black strain. We compared the black-substituted average fractions *F_i_* (the dashed lines) to the real fractions as a function of radius (the solid lines). At a small radius relative to *R*_0_ = 3.5 mm, the error from introducing a black strain was large; this is likely because we defined black as the absence of any other channels and channels typically had large overlaps close to the homeland. At large radius, the error from introducing a black strain was negligible.

**Fig S2.**
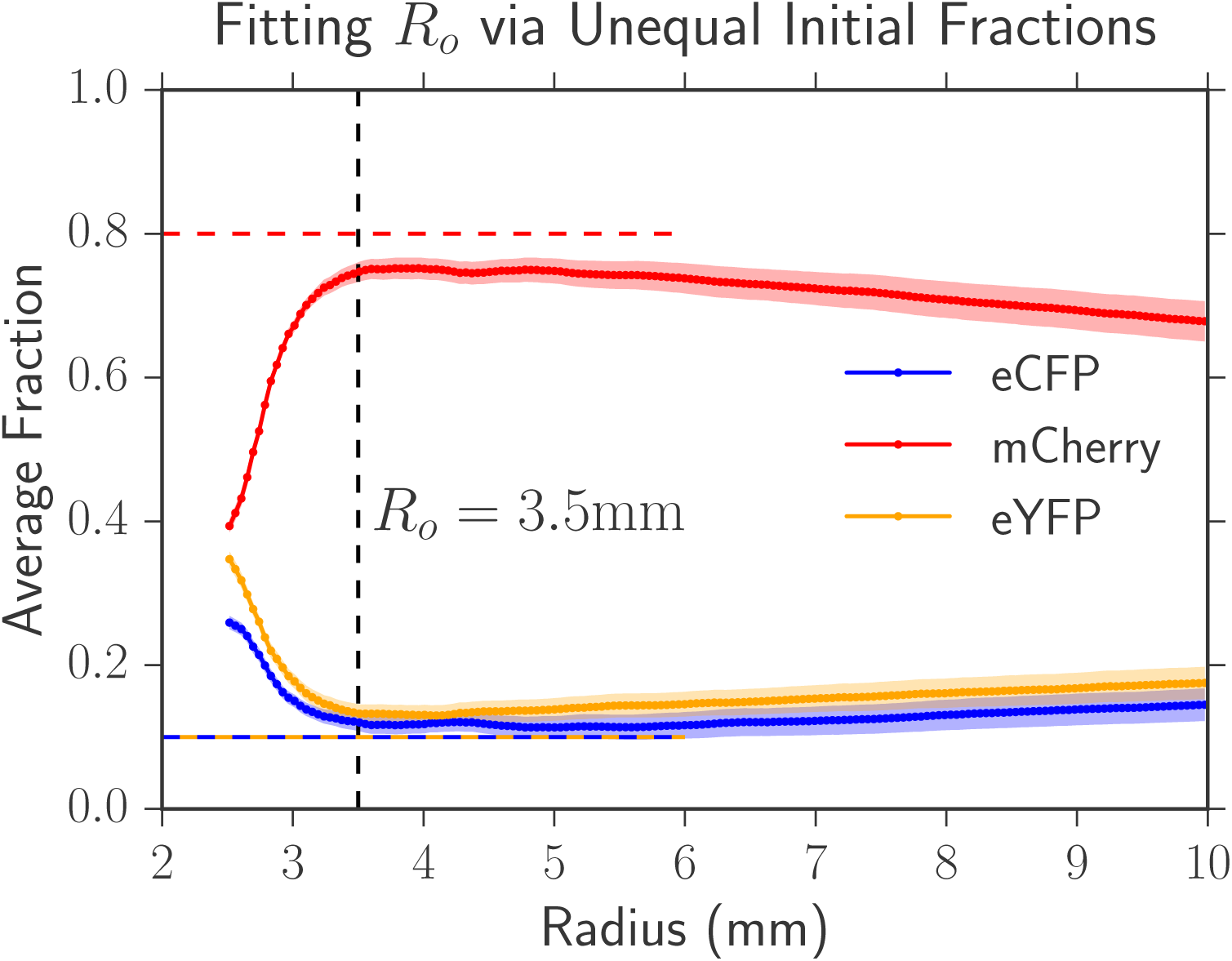
Determining *R*_0_. To fit the radius *R*_0_ where our image analysis package became accurate, we inoculated 80% of mCherry, 10% of eCFP, and 10% of eYFP in 10 range expansions and tabulated the average fraction of each strain. The inoculated fractions are illustrated by dashed lines. As seen in the plot, at a radius of approximately *R*_0_ = 3.50 ± 0.05 mm the average fractions approximately matched the initial inoculated fractions. Our image analysis package inaccurately predicted fractions in the homeland because of significant overlap between the strains.

**Fig S3.**
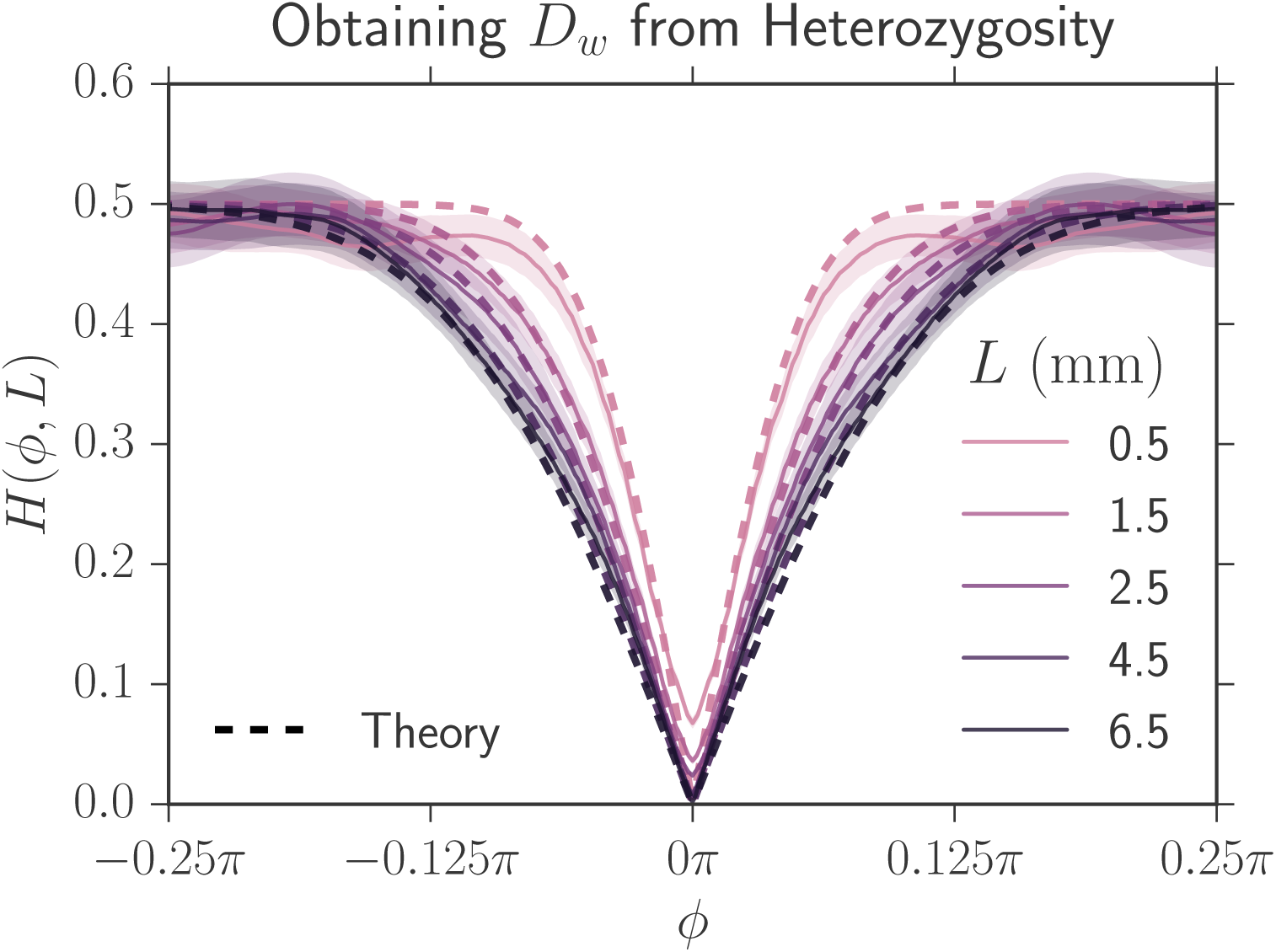
Error bars when fitting *D_w_*. The same as the right side of Figure 5 except with error bars; the shaded areas are the standard error of the mean.

**Fig S4.**
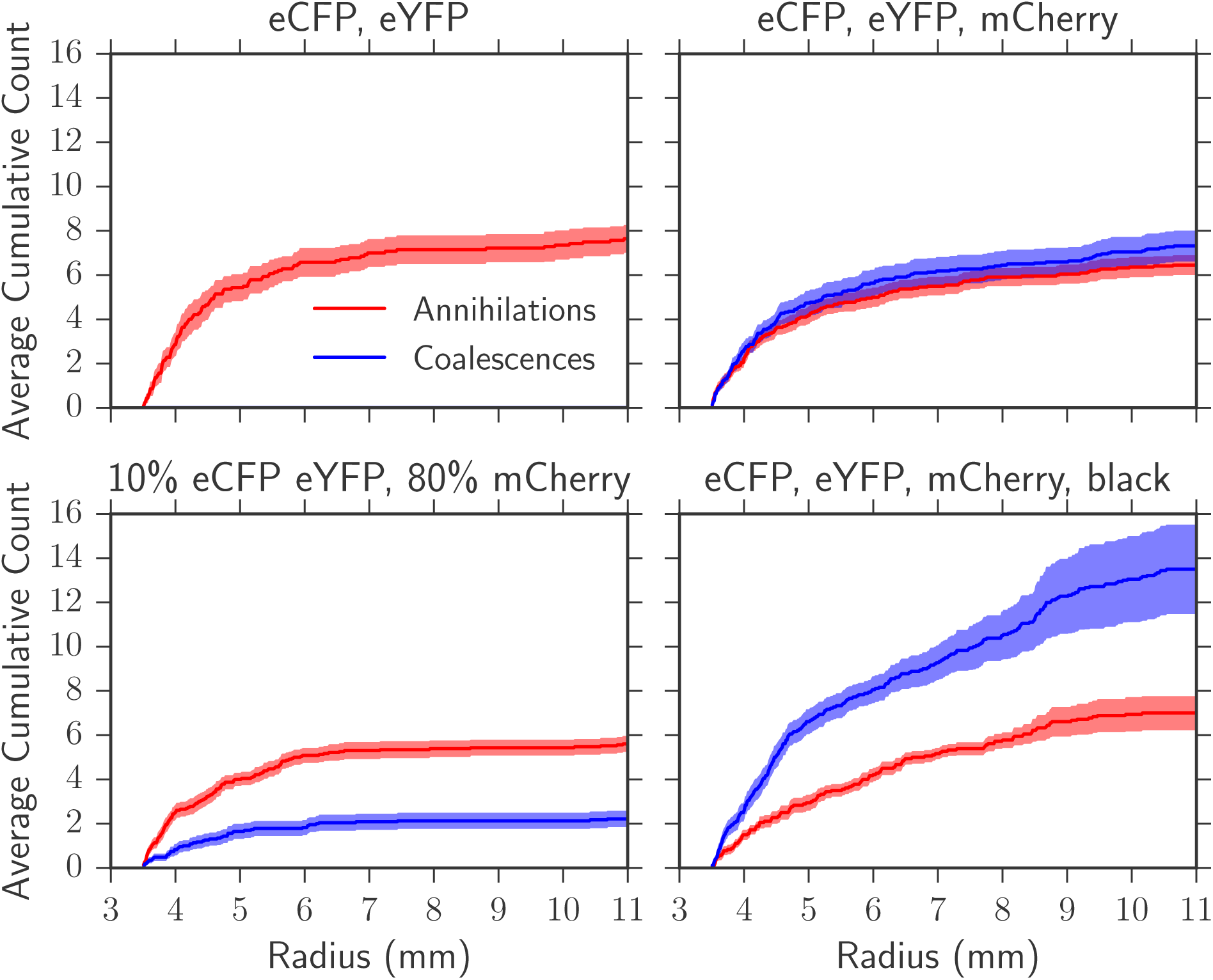
Average cumulative annihilations and coalescences for two, three, and four strains. All strains were inoculated in equal fractions except for the experiment with 10% of eCFP, 10% of eYFP and 80% of mCherry. The annihilation and coalescence rates (the slope of the respective curves) decrease as radius increases as there are less domain walls due to previous collisions and also because inflation decreases the probability of two walls colliding per length expanded. As the number of colors increases, coalescences occur more often than annihilations.

**Fig S5.**
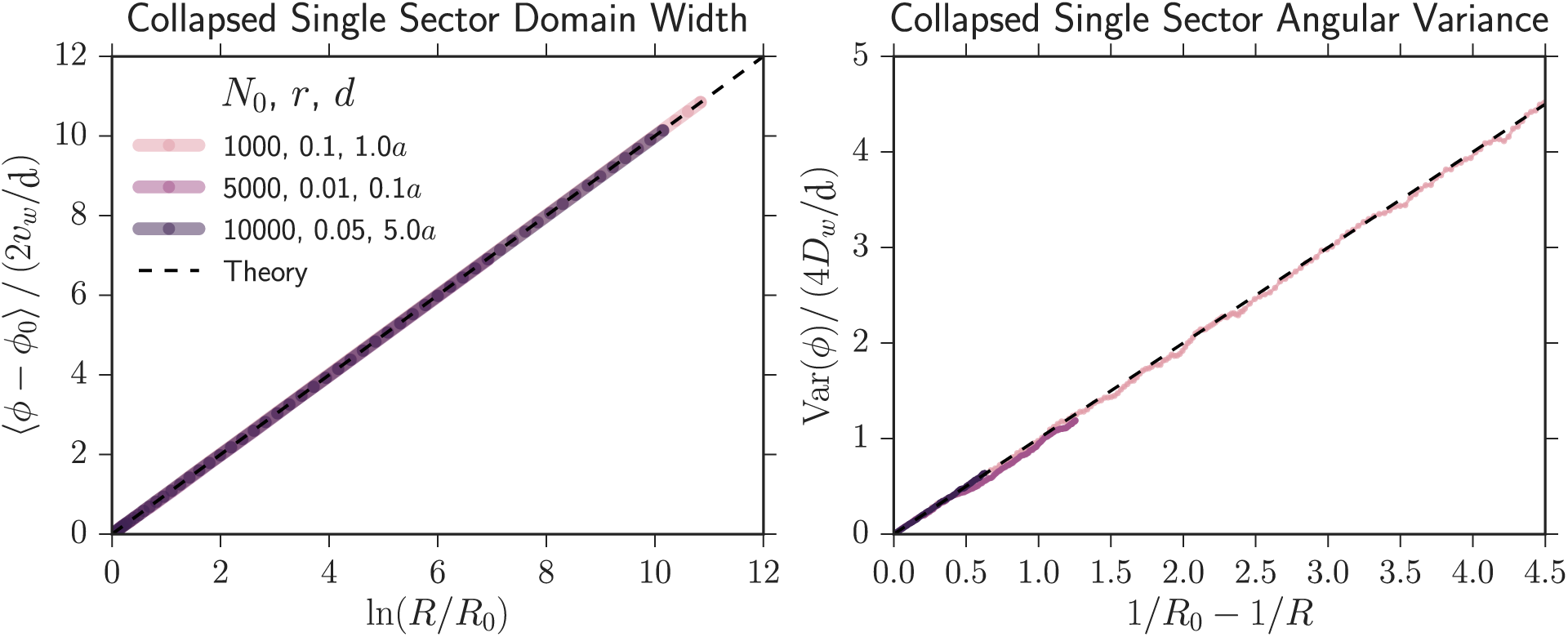
Confirming simulation accuracy. We simulated a single fit sector sweeping through a less fit strain. It is expected that the fit strain sector dynamics satisfy 〈*ϕ* − *ϕ*_0_〉 = 2*v_w_* ln (*R*/*R*_0_) and Var (*ϕ*) = 4*D_w_* (1/*R*_0_ − 1/*R*), as seen in Appendix S1. To test that our simulation appropriately reproduced this behavior, we quantified the average angular growth 〈*ϕ* − *ϕ*_0_〉 and angular variance Var(*ϕ*) as we varied the simulation parameters *N*_0_ (initial number of cells), *r* (selective advantage of the fitter strain), and *d* (distance the colony expanded each generation). The cell width, *a*, was kept constant. These parameters relate to the sector dynamics via *D_w_* = *a*^2^/(2*d*), 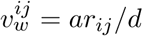, and *R*_0_ = (*N*_0_*a*)/(2*π*). We confirmed that both the average angular growth 〈*ϕ* − *ϕ*_0_〉 and angular variance Var(*ϕ*) had the correct functional form and dependence on the microscopic parameters (the dashed black line). In the main text, we used *d* = *a* for simplicity.

**Fig S6.**
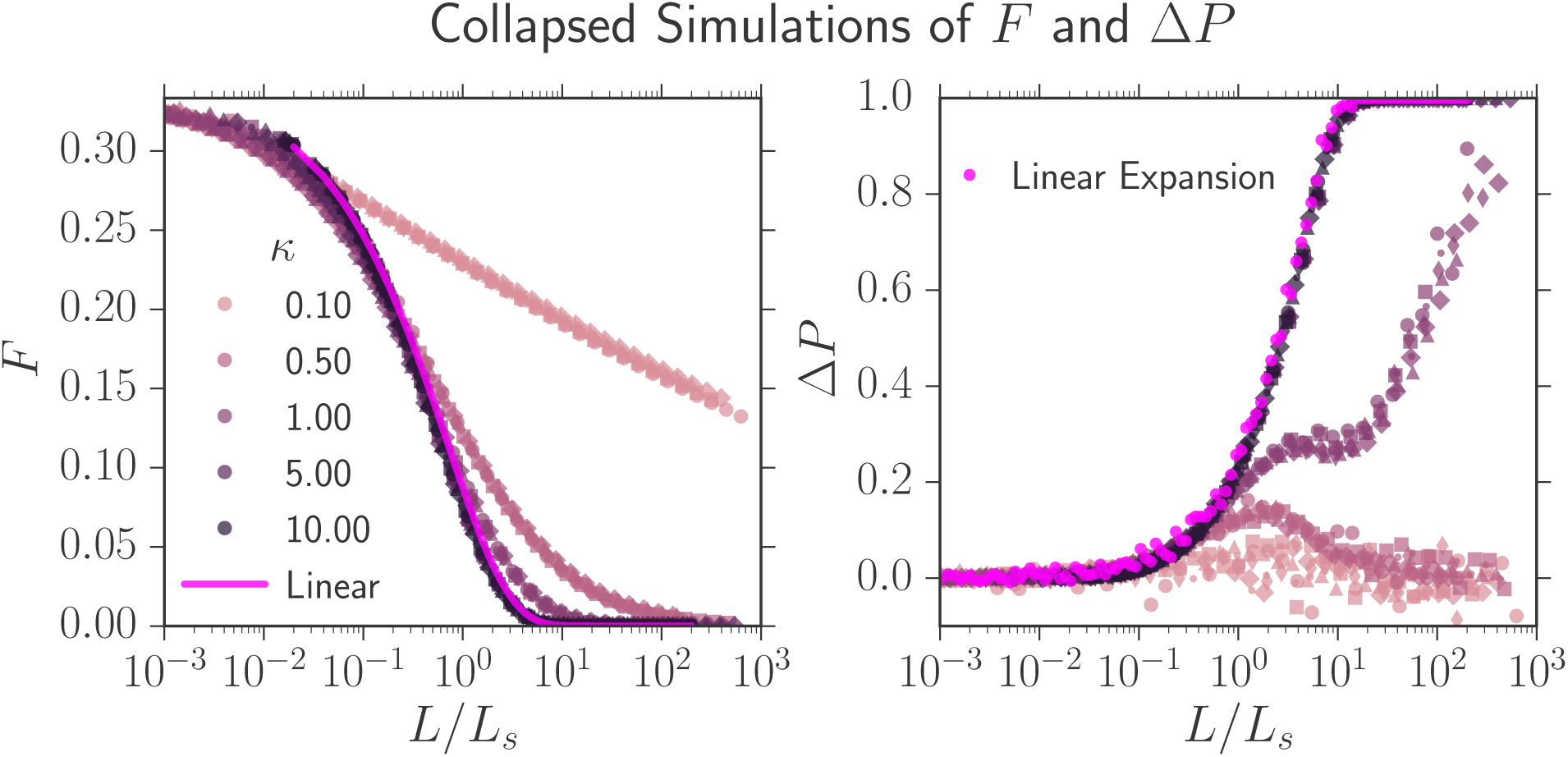
Collapsed average fraction and annihilation asymmetry on a linear scale. Identical to Figure 11 except the *y*-axis of *F*(*L*/*L_s_*, *κ*) is placed on a linear scale, which may be useful for comparison with experiments.

**Fig S7.**
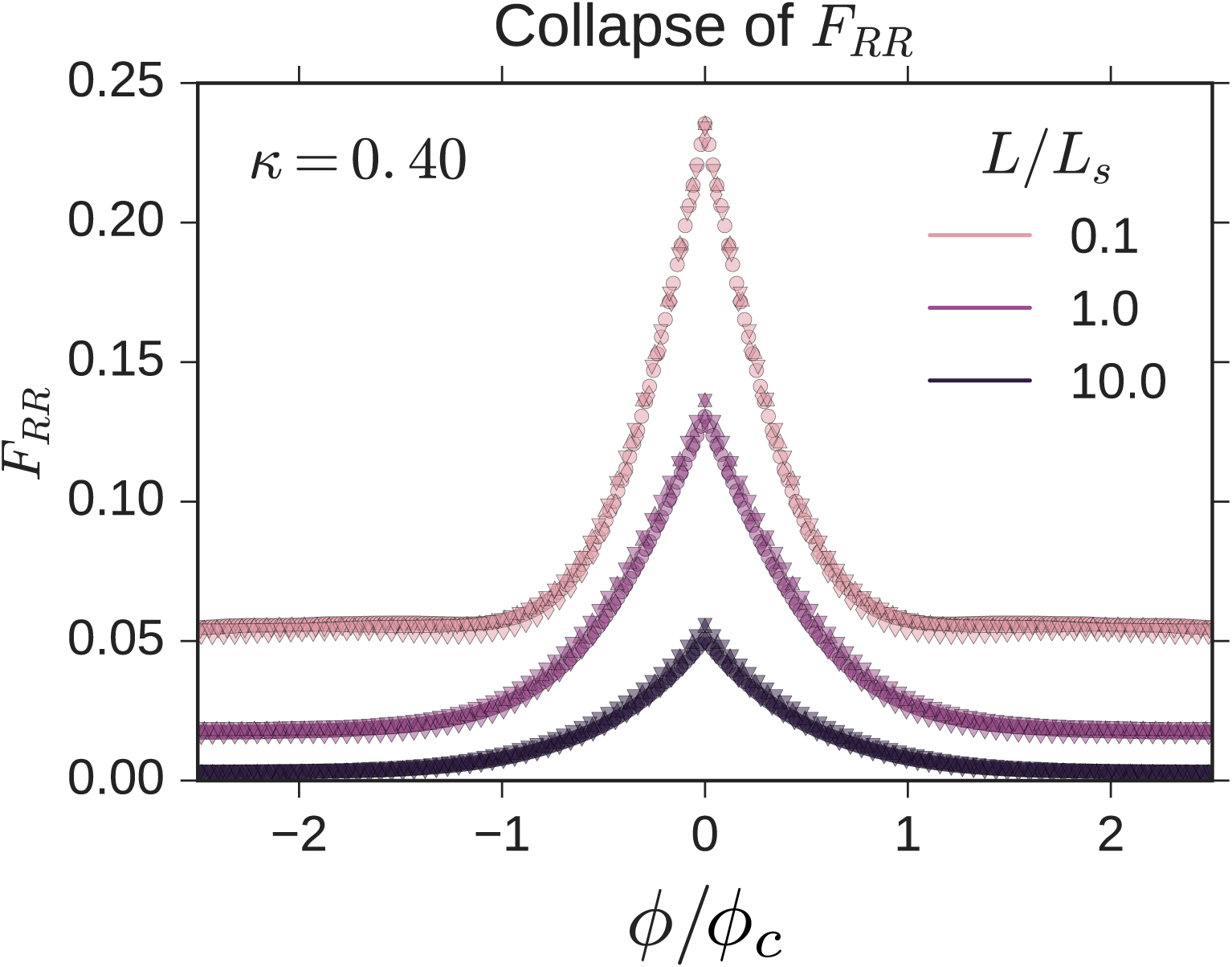
Collapse of *F_ij_*. We ran four simulations where we varied *v_w_* (the velocity that the two other more fit strains swept through the less fit strain) and *R*_0_ such that 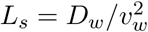 changed but 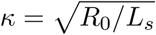 was fixed. Each simulation has a different symbol in the plot. We found that *F_ij_* could be collapsed at the same *L*/*L_s_* as long as *κ* remained fixed (we arbitrarily set it to *κ* = 0.4) and as long as the angular variable *ϕ* was rescaled by 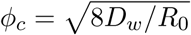. If *ϕ_c_* approached the system size *ϕ_c_* ≈ 2*π*, *F_ij_* could not be collapsed onto the above curves due to finite size effects. Note that even though we only show *F_RR_*, all correlation functions *F_ij_* could be collapsed using this procedure.

**Fig S8.**
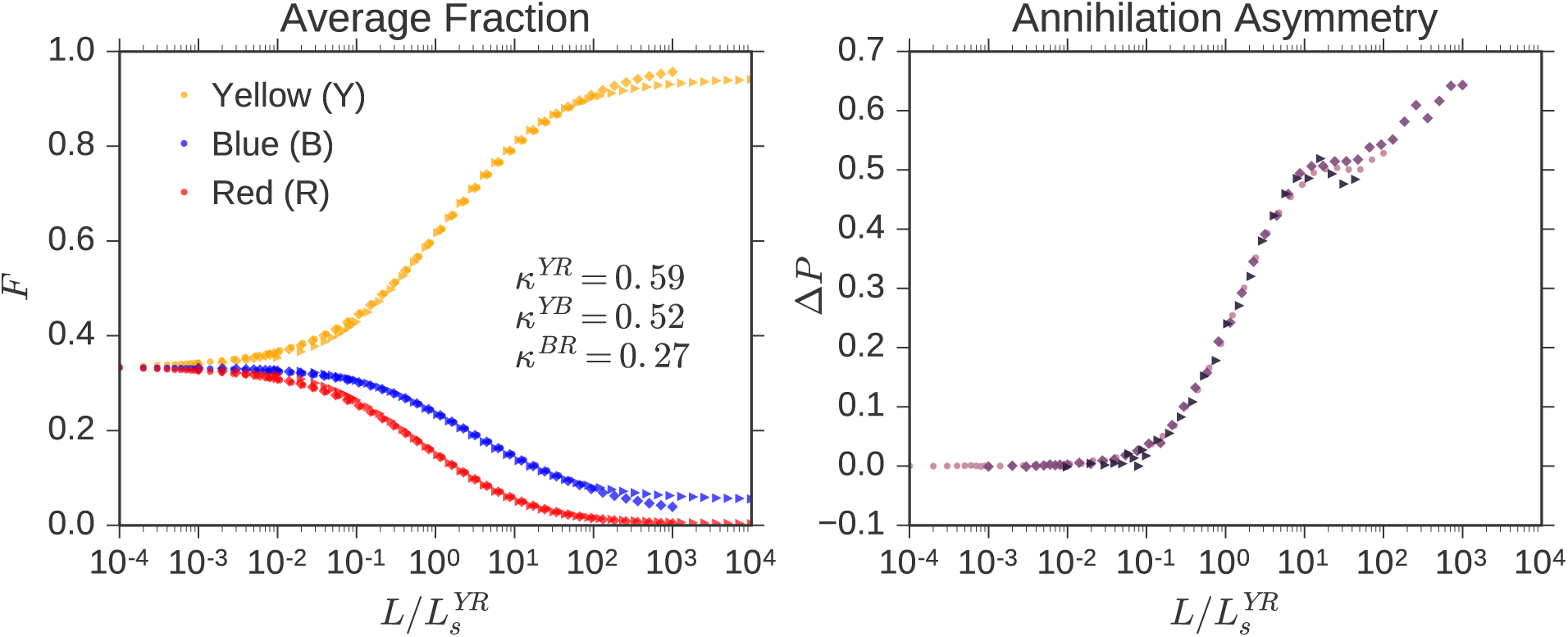
Collapse of average fraction and annihilation asymmetry. We simulated three competing strains and arbitrarily chose a fixed set of 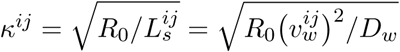 to reflect differing selective advantages between the strains. We set the initial radius of the expansion *R*_0_ = *N*_0_*a*/(2*π*) to *N*_0_ = {200, 2000, 20000} in three simulations and altered *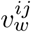* to tune the set of *κ^ij^*’s to their fixed values. We found that the dynamics collapsed as long as *L* was rescaled by any selection length scale in the system, i.e. *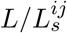* (we chose to use *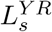*). The diamond simulation, corresponding to *N*_0_ = 200, deviated slightly from the other simulations because of finite size effects (i.e. when *ϕ_c_* ~ 2*π*).

## References

1. Lang GI, Botstein D, Desai MM. Genetic Variation and the Fate of Beneficial Mutations in Asexual Populations. Genetics. 2011;188(3):647–661. doi:10.1534/genetics.111.128942.

2. Korolev KS, Avlund M, Hallatschek O, Nelson DR. Genetic demixing and evolution in linear stepping stone models. Reviews of Modern Physics. 2010;82(2):1691–1718. doi:10.1103/RevModPhys.82.1691.

3. Nadell CD, Drescher K, Foster KR. Spatial structure, cooperation and competition in biofilms. Nature Reviews Microbiology. 2016;14(9):589–600. doi:10.1038/nrmicro.2016.84.

4. Sprouffske K, Merlo LMF, Gerrish PJ, Maley CC, Sniegowski PD. Cancer in Light of Experimental Evolution. Current Biology. 2012;22(17):R762–R771. doi:10.1016/j.cub.2012.06.065.

5. Langin KM, Sillett TS, Funk WC, Morrison SA, Desrosiers MA, Ghalambor CK. Islands within an island: Repeated adaptive divergence in a single population. Evolution. 2015;69(3):653–665. doi:10.1111/evo.12610.

6. Donaldson GP, Lee SM, Mazmanian SK. Gut biogeography of the bacterial microbiota. Nature Reviews Microbiology. 2015;14(1):20–32. doi:10.1038/nrmicro3552.

7. Morrison SJ, Spradling AC. Stem Cells and Niches: Mechanisms That Promote Stem Cell Maintenance throughout Life. Cell. 2008;132(4):598–611. doi:10.1016/j.cell.2008.01.038.

8. Hallatschek O, Hersen P, Ramanathan S, Nelson DR. Genetic drift at expanding frontiers promotes gene segregation. Proceedings of the National Academy of Sciences. 2007;104(50):19926–19930. doi:10.1073/pnas.0710150104.

9. Gralka M, Stiewe F, Farrell F, Möbius W, Waclaw B, Hallatschek O. Allele surfing promotes microbial adaptation from standing variation. Ecology Letters. 2016;19(8):889–898. doi:10.1111/ele.12625.

10. Lavrentovich MO, Koschwanez JH, Nelson DR. Nutrient shielding in clusters of cells. Physical Review E. 2013;87(6):062703. doi:10.1103/PhysRevE.87.062703.

11. Hartl DL, Clark AG. Principles of Population Genetics. 4th ed. Sunderland, MA: Sinauer Associates, Inc.; 2007. Available from: http://www.sinauer.com/principles-of-population-genetics.html.

12. White Ta, Perkins SE, Heckel G, Searle JB. Adaptive evolution during an ongoing range expansion: the invasive bank vole (Myodes glareolus) in Ireland. Molecular Ecology. 2013;22(11):2971–2985. doi:10.1111/mec.12343.

13. Phillips BL, Brown GP, Webb JK, Shine R. Invasion and the evolution of speed in toads. Nature. 2006;439(7078):803–803. doi:10.1038/439803a.

14. Templeton A. Out of Africa again and again. Nature. 2002;416(6876):45–51. doi:10.1038/416045a.

15. Gerrish PJ, Lenski RE. The fate of competing beneficial mutations in an asexual population. Genetica. 1998;102-103(1-6):127–44. doi:10.1023/A:1017067816551.

16. Baym M, Lieberman TD, Kelsic ED, Chait R, Gross R, Yelin I, et al. Spatiotemporal microbial evolution on antibiotic landscapes. Science. 2016;353(6304):1147–1151. doi:10.1126/science.aag0822.

17. Korolev KS, Müller MJI, Karahan N, Murray AW, Hallatschek O, Nelson DR. Selective sweeps in growing microbial colonies. Physical Biology. 2012;9(2):026008. doi:10.1088/1478-3975/9/2/026008.

18. Korolev KS, Xavier JB, Nelson DR, Foster KR. A Quantitative Test of Population Genetics Using Spatiogenetic Patterns in Bacterial Colonies. The American Naturalist. 2011;178(4):538–552. doi:10.1086/661897.

19. Doering CR, Ben-Avraham D. Interparticle distribution functions and rate equations for diffusion-limited reactions. Physical Review A. 1988;38(6):3035–3042. doi:10.1103/PhysRevA.38.3035.

20. Amar JG, Family F. Diffusion annihilation in one dimension and kinetics of the Ising model at zero temperature. Physical Review A. 1990;41(6):3258–3262. doi:10.1103/PhysRevA.41.3258.

21. Krapivsky PL, Redner S, Ben-Naim E. A Kinetic View of Statistical Physics. Cambridge: Cambridge University Press; 2010. Available from: http://ebooks.cambridge.org/ref/id/CBO9780511780516.

22. Potts RB, Domb C. Some generalized order-disorder transformations. Mathematical Proceedings of the Cambridge Philosophical Society. 1952;48(October):106. doi:10.1017/S0305004100027419.

23. Wu FY. The Potts model. Reviews of Modern Physics. 1982;54(I):235–268. doi:10.1103/RevModPhys.54.235.

24. Glauber RJ. Time-Dependent Statistics of the Ising Model. Journal of Mathematical Physics. 1963;4(2):294. doi:10.1063/1.1703954.

25. Clifford P, Sudbury A. A model for spatial conflict. Biometrika. 1973;60(3):581–588. doi:10.1093/biomet/60.3.581.

26. Ódor G. Universality classes in nonequilibrium lattice systems. Reviews of Modern Physics. 2004;76(3):663–724. doi:10.1103/RevModPhys.76.663.

27. Masser T, Ben-Avraham D. Kinetics of coalescence, annihilation, and the q-state Potts model in one dimension. Physics Letters A. 2000;275(5-6):382–385. doi:10.1016/S0375-9601(00)00622-8.

28. Masser TO, Ben-Avraham D. Correlation functions for diffusion-limited annihilation, *A* + *A* → *A*. Physical Review E. 2001;64(6):062101. doi:10.1103/PhysRevE.64.062101.

29. Derrida B, Zeitak R. Distribution of domain sizes in the zero temperature Glauber dynamics of the one-dimensional Potts model. Physical Review E. 1996;54(3):2513–2525. doi:10.1103/PhysRevE.54.2513.

30. Ali A, Grosskinsky S. Pattern formation through genetic drift at expanding population fronts. Advances in Complex Systems. 2009;13(03):349–366. doi:10.1142/S0219525910002578.

31. Ali A, Ball RC, Grosskinsky S, Somfai E. Scale-invariant growth processes in expanding space. Physical Review E. 2013;87(2):020102. doi:10.1103/PhysRevE.87.020102.

32. Lavrentovich MO, Korolev KS, Nelson DR. Radial Domany-Kinzel models with mutation and selection. Physical Review E. 2013;87(1). doi:10.1103/PhysRevE.87.012103.

33. Sen P, Ray P. *A* + *A* → ∅ model with a bias towards nearest neighbor. Physical Review E. 2015;92(1):012109. doi:10.1103/PhysRevE.92.012109.

34. Weinstein B. Range Expansions on GitHub; 2016. Available from: https://github.com/Range–Expansions.

35. Weber MF, Poxleitner G, Hebisch E, Frey E, Opitz M. Chemical warfare and survival strategies in bacterial range expansions. Journal of The Royal Society Interface. 2014;11(96):20140172–20140172. doi:10.1098/rsif.2014.0172.

36. Jauffred L, Munk Vejborg R, Korolev KS, Brown S, Oddershede LB. Chirality in microbial biofilms is mediated by close interactions between the cell surface and the substratum. The ISME Journal. 2017;x:xx–xx. doi:10.1038/ismej.2017.19.

37. Amann E, Ochs B, Abel KJ. Tightly regulated tac promoter vectors useful for the expression of unfused and fused proteins in Escherichia coli. Gene. 1988;69(2):301–315. doi:10.1016/0378-1119(88)90440-4.

38. Möbius W, Murray AW, Nelson DR. How Obstacles Perturb Population Fronts and Alter Their Genetic Structure. PLOS Computational Biology. 2015;11(12):e1004615. doi:10.1371/journal.pcbi.1004615.

39. Zuiderveld K. Contrast limited adaptive histogram equalization. Graphics Gems IV. 1994; p. 474–485.

40. Schindelin J, Arganda-Carreras I, Frise E, Kaynig V, Longair M, Pietzsch T, et al. Fiji: an open-source platform for biological-image analysis. Nature Methods. 2012;9(7):676–682. doi:10.1038/nmeth.2019.

41. Bennett CH. Serially Deposited Amorphous Aggregates of Hard Spheres. Journal of Applied Physics. 1972;43(6):2727. doi:10.1063/1.1661585.

42. Lavrentovich MO, Nelson DR. Survival probabilities at spherical frontiers. Theoretical Population Biology. 2015;102:26–39. doi:10.1016/j.tpb.2015.03.002.

43. Harper M. Python-ternary: A python library for ternary plots; 2011. Available from: https://github.com/marcharper/python-ternary.

## References

1. Korolev KS, Avlund M, Hallatschek O, Nelson DR. Genetic demixing and evolution in linear stepping stone models. Reviews of Modern Physics. 2010;82(2):1691–1718. doi:10.1103/RevModPhys.82.1691.

2. Masser TO, Ben-Avraham D. Correlation functions for diffusion-limited annihilation, *A* + *A* → *A*. Physical Review E. 2001;64(6):062101. doi:10.1103/PhysRevE.64.062101.

3. Masser T, Ben–Avraham D. Kinetics of coalescence, annihilation, and the q–state Potts model in one dimension. Physics Letters A. 2000;275(5-6):382–385. doi:10.1016/S0375-9601(00)00622-8.

4. Derrida B, Zeitak R. Distribution of domain sizes in the zero temperature Glauber dynamics of the one-dimensional Potts model. Physical Review E. 1996;54(3):2513–2525. doi:10.1103/PhysRevE.54.2513.

5. Korolev KS, Xavier JB, Nelson DR, Foster KR. A Quantitative Test of Population Genetics Using Spatiogenetic Patterns in Bacterial Colonies. The American Naturalist. 2011;178(4):538–552. doi:10.1086/661897.

6. Lavrentovich MO, Korolev KS, Nelson DR. Radial Domany-Kinzel models with mutation and selection. Physical Review E. 2013;87(1). doi:10.1103/PhysRevE.87.012103.

7. Kimura M, Weiss GH. The Stepping Stone Model of Population Structure and the Decrease of Genetic Correlation with Distance. Genetics. 1964;49(4):561–576. doi:10.1093/oxfordjournals.molbev.a025590.

8. Potts RB, Domb C. Some generalized order-disorder transformations. Mathematical Proceedings of the Cambridge Philosophical Society. 1952;48(October):106. doi:10.1017/S0305004100027419.

9. Wu FY. The Potts model. Reviews of Modern Physics. 1982;54(I):235–268. doi:10.1103/RevModPhys.54.235.

10. Glauber RJ. Time-Dependent Statistics of the Ising Model. Journal of Mathematical Physics. 1963;4(2):294. doi:10.1063/1.1703954.

11. Moran PAP. Random processes in genetics. Mathematical Proceedings of the Cambridge Philosophical Society. 1958;54(01):60–71. doi:10.1017/S0305004100033193.

12. Good BH, Desai MM. Fluctuations in fitness distributions and the effects of weak linked selection on sequence evolution. Theoretical Population Biology. 2013;85:86–102. doi:10.1016/j.tpb.2013.01.005.

13. Gardiner C. Stochastic Methods: A Handbook for the Natural and Social Sciences. 4th ed. Springer; 2009. Available from: http://www.amazon.com/Stochastic–Methods–Handbook–Sciences–Synergetics/dp/3540707123.

14. Ali A, Grosskinsky S. Pattern formation through genetic drift at expanding population fronts. Advances in Complex Systems. 2009;13(03):349–366. doi:10.1142/S0219525910002578.

15. Ali A, Ball RC, Grosskinsky S, Somfai E. Scale-invariant growth processes in expanding space. Physical Review E. 2013;87(2):020102. doi:10.1103/PhysRevE.87.020102.

16. Hallatschek O, Nelson DR. Life at the front of an expanding population. Evolution; international journal of organic evolution. 2010;64(1):193–206. doi:10.1111/j.1558-5646.2009.00809.x.

18. Gralka M, Stiewe F, Farrell F, Möbius W, Waclaw B, Hallatschek O. Allele surfing promotes microbial adaptation from standing variation. Ecology Letters. 2016;19(8):889–898. doi:10.1111/ele.12625.

## References

1. Korolev KS, Müller MJI, Karahan N, Murray AW, Hallatschek O, Nelson DR. Selective sweeps in growing microbial colonies. Physical Biology. 2012;9(2):026008. doi:10.1088/1478-3975/9/2/026008.

2. Gralka M, Stiewe F, Farrell F, Möbius W, Waclaw B, Hallatschek O. Allele surfing promotes microbial adaptation from standing variation. Ecology Letters. 2016;19(8):889–898. doi:10.1111/ele.12625.

